# Order and disorder – an integrative structure of the full-length human growth hormone receptor

**DOI:** 10.1101/2020.06.25.171116

**Authors:** Noah Kassem, Raul Araya-Secchi, Katrine Bugge, Abigail Barclay, Helena Steinocher, Adree Khondker, Aneta J. Lenard, Jochen Bürck, Anne S. Ulrich, Martin Cramer Pedersen, Yong Wang, Maikel C. Rheinstädter, Per Amstrup Pedersen, Kresten Lindorff-Larsen, Lise Arleth, Birthe B. Kragelund

## Abstract

Despite the many physiological and pathophysiological functions of the human growth hormone receptor (hGHR), a detailed understanding of its *modus operandi* is hindered by the lack of structural information of the entire receptor at the molecular level. Due to its relatively small size (70 kDa) and large content of structural disorder (>50%), this membrane protein falls between the cracks of conventional high-resolution structural biology methods. Here, we study the structure of the full-length hGHR in nanodiscs with small angle-X-ray scattering (SAXS) as the foundation. We developed an approach in which we combined SAXS, X-ray diffraction and NMR spectroscopy obtained on the individual domains and integrated the data through molecular dynamics simulations to interpret SAXS data on the full-length hGHR in nanodiscs. The structure of the hGHR was determined in its monomeric state and provides the first experimental model of any full-length cytokine receptor in a lipid membrane. Combined, our results highlight that the three domains of the hGHR are free to reorient relative to each other, resulting in a broad structural ensemble. Our work exemplifies how integrating experimental data from several techniques computationally, may enable the characterization of otherwise inaccessible structures of membrane proteins with long disordered regions, a widespread phenomenon in biology. To understand orchestration of cellular signaling by disordered chains, the hGHR is archetypal and its structure emphasizes that we need to take a much broader, ensemble view on signaling.

## INTRODUCTION

The human growth hormone receptor (hGHR) is ubiquitously expressed^1^, and is activated by human growth hormone (hGH), produced in the pituitary gland. hGHR is important for regulating growth at a cellular and systemic level^1,2^, and is involved in the regulation of hepatic metabolism, cardiac function, bone turnover and the immune system^3^. Besides direct promotion of growth^4^, its ligand hGH can also indirectly regulate growth by initiating the synthesis of insulin-like growth factor-I (IGF-I), an important factor in postnatal growth^2,5,6^. Excess hGH production and mutations in the hGHR gene manifest in different diseases including cancer^7^ and growth deficiencies^8–11^, with associated cardiovascular, metabolic and respiratory difficulties^8^, and both hGH-based agonists and antagonists of the receptor exist as approved drugs^12,13^.

The hGHR is one of ~40 receptors belonging to the class 1 cytokine receptor family^14^. The family is topologically similar with a tripartite structure consisting of a folded extracellular domain (ECD), a single-pass transmembrane domain (TMD), and a disordered intracellular domain (ICD)^14–16^. A characteristic trait of these receptors is the lack of intrinsic kinase activity, with the ICD instead forming a binding platform for a variety of signaling kinases and regulatory proteins^15,17,18^, as well as of certain specific membrane lipids^16^ (**Fig. 1A)**. Within the ECD, the receptors share a characteristic cytokine receptor homology domain consisting of two fibronectin type III domains (D1, N-terminal and D2, C-terminal), each with a seven stranded β-sandwich structure. Two hallmark disulfide bonds and a conserved WSXWS motif (X is any amino acid)^19,20^ located in D1 and D2, respectively, are suggested to be important for cell surface localization and discrimination between signaling pathways^19,21^. In hGHR, this motif is instead YGEFS^17^, but the reason for this variation has remained enigmatic. Beside hGHR, group 1 of the class 1 cytokine receptor also encompasses the prolactin receptor (PRLR) and the erythropoietin (EPO) receptor. This group is considered to be the most structurally simple with one cytokine receptor homology domain and ligand binding in a 2:1 complex^17,18^.

**Figure 1.**
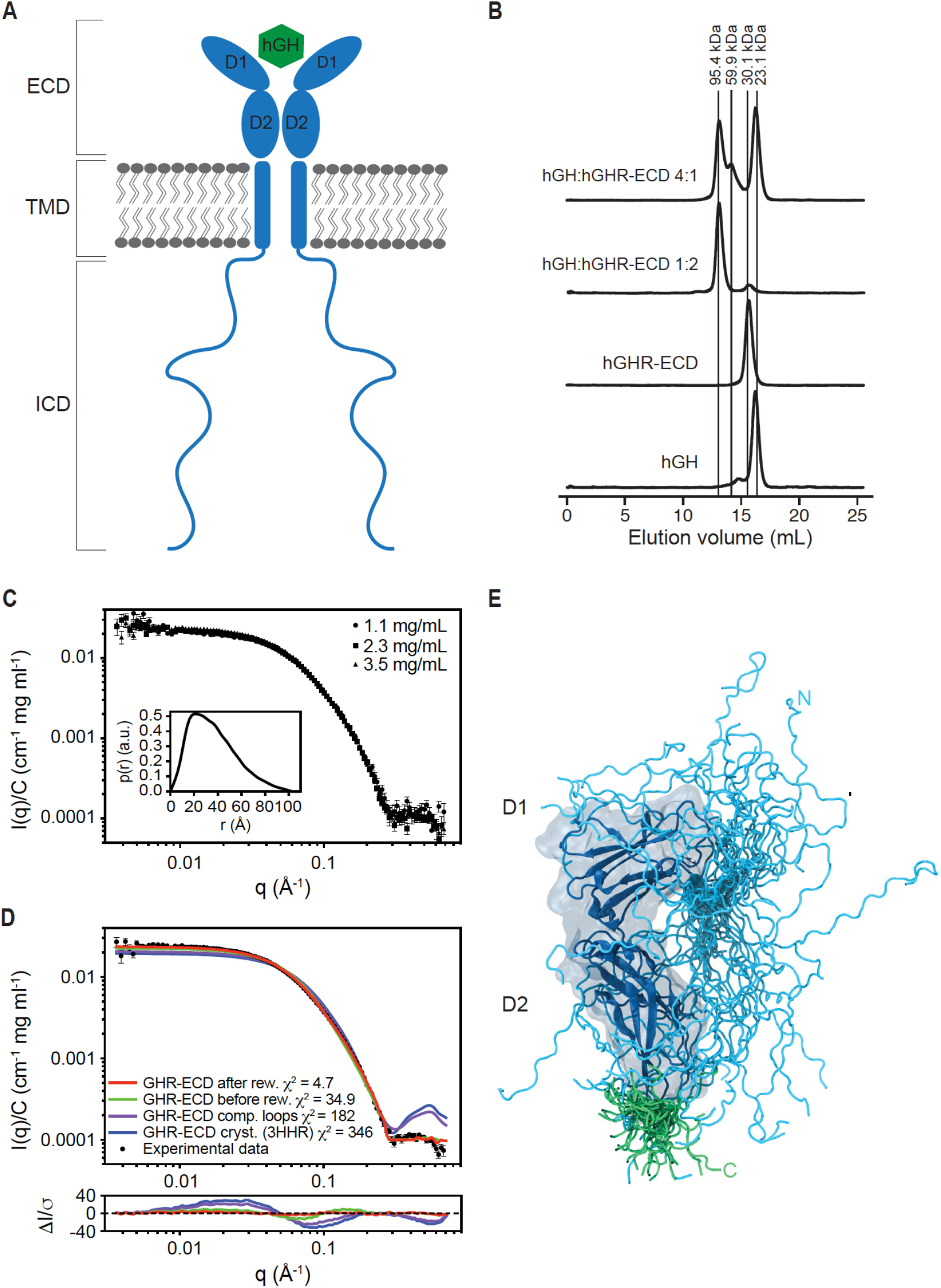
The hGHR has a dynamic ECD with a broad structural ensemble. (A) A schematic representation of homodimeric hGHR (blue) in the membrane in complex with hGH (green). ECD, Extracellular domain; TMD, transmembrane domain and ICD, intracellular domain. (B) SEC profiles of hGHR-ECD and hGH in 20 mM Na_2_HPO_4_/NaH_2_PO_4_ (pH 7.4), 150 mM NaCl at ratios 1:0 (hGH:hGHRECD 1:0), 0:1 (hGH:hGHR-ECD 0:1), 1:2 (hGH:hGHR-ECD 1:2), 4:1 (hGH:hGHR-ECD 4:1). Absorption was measured at 280 nm. (C) Concentration-normalized SAXS data from hGHR-ECD (concentrations in legend) with the *p(r)* from the 3.5 mg/mL sample shown as insert. (D) SAXS data from hGHR-ECD at 3.5 mg/ml (black dots) together with fits of the theoretical scattering curves from a crystal structure of hGRH-ECD (blue, PDB 3HHR), the same crystal structure with missing loops completed (purple), and the average (green), and reweighted average (red) of scattering curves of the 500 hGHR-ECD models with added N- and C-terminal tails. Residuals are plotted below. (E) An ensemble model of the hGHR-ECD with a representative reweighted sub-ensemble of 100 models highlighting the N- (cyan) and C- (green) terminal dynamic tails.

Receptor activation is achieved by hGH binding to hGHR via two asymmetric binding sites^22^, leading to structural rearrangements that are propagated through the TMD to the ICD^23^. A recent study found that when hGH binds to a pre-formed hGHR dimer, structural rearrangements in the ECD leads to separation of the ICDs just below the TMD^23^. This leads to activation through cross-phosphorylation of the Janus kinases 2 (JAK2) bound at the proline rich Box1-motif in the juxtamembrane region^23^. Furthermore, this study demonstrated that receptor dimerization in isolation is insufficient for receptor activation^23^. Nonetheless, while recent single-particle tracking studies suggested dimerization to depend on expression levels^24^, it is still debated to what extent the hGHR exists as pre-formed dimers *in vivo*^25^, or if the hGHR only dimerizes upon hGH binding^26^.

From the viewpoint of structural biology, the hGH/hGHR system has a high molecular complexity with ordered and disordered domains joined by a minimal membrane embedded part. Hence, structural characterization of this receptor has so far utilized a divide and conquer approach, where the domains have been studied in isolation. This includes the crystal structures of the ECD in the monomeric state^25^, in 1:1^27^- and 2:1^22^ complexes with hGH, and of hGH alone^28^. Furthermore, structures of the dimeric state of the hGHR-TMD in detergent micelles have been solved by nuclear magnetic resonance (NMR)^29^ spectroscopy, while the hGHR-ICD was shown by NMR to adopt a fully intrinsically disordered region (IDR)^16^. A recent approach that combined experimental data with computational efforts provided a model of the similar PRLR monomer built from integration of several individual sets of experimental data recorded on isolated domains^30^. This work provided the first view of a full-length class 1 cytokine receptor. However, no structure or model based on data collected on an intact, full-length class I cytokine receptor exists, leaving a blind spot for how the domains effect each other and are spatially organized.

Even with the major advances in cryo-electron microscopy (EM)^31^, the full-length hGHR remains a challenge to structural biology. With 70 kDa, the receptor is a small target for cryo-EM, but adding to this, the fact that more than 50% of the protein is intrinsically disordered leaves only ~30 kDa visible. Likewise, the intrinsic disorder of the ICD also hampers crystallographic studies. Orthogonally, 70 kDa plus membrane mimetics makes up too large a target for NMR, where the combined molecular properties would lead to slow tumbling and severe line broadening. Hence, the hGHR appears to be an orphan to structural biology, along with a large group of other membrane proteins with long, disordered regions, including most of the ~1400 human single pass membrane proteins^32^. Lower resolution techniques, such as solution small-angle X-ray- and neutron scattering (SAXS/SANS) offer important alternatives and provide information about the solution structure of a protein regardless of whether it is disordered or not. These methods become particularly strong in combination with experimental information from orthogonal techniques through computational modelling. In such situations, SAS data allow for refining a low-resolution structure of a protein, including membrane proteins^33,34^. Recent advances building on the use of nanodiscs^35^, have further proved the applicability of SAS in membrane protein structural biology when combined with computational modeling^36,37^. Remarkably however, no membrane protein with the degree of disorder seen in hGHR has previously been studied in a nanodisc or approached by SAS.

Here we applied an integrative approach to access the structure of the monomeric hGHR from SAXS data recorded on the full-length receptor in a nanodisc. The data were validated and interpreted by combining SAXS, NMR and X-ray diffraction (XRD) data obtained on the individual domains of hGHR through computational modeling. This has resulted in the first experimentally supported structure model based on studies of an intact, full-length, single-pass cytokine receptor in a lipid membrane; a topology which represents ~40 human cytokine receptors and many other membrane proteins. Our approach exemplifies that combining SAS and computational modeling could be the bridge required for accessing structural information on the ~1400 single-pass receptors in humans^38^.

## RESULTS

To arrive at the final result of this work we took on a three-step approach. First, to aid the analysis of SAS data on the full-length hGHR and qualify the integrity of the methodology, several different biophysical data were acquired and analyzed on isolated, individual parts of the hGHR. Secondly, SAS data were acquired on the full-length hGHR in nanodiscs, expressed in yeast cells and carrying a C-terminal GFP-deca-histidine tag (GFP-H_10_). Finally, all the data were interpreted and integrated using molecular dynamics simulations.

### The binding competent hGHR-ECD solution state ensemble contains disorder

While crystal structures of an N- and C-terminal truncated version of the hGHR-ECD exist^22,27,28^, the complete hGHR-ECD has not previously been studied in solution. Therefore, to describe the ensemble of the full domain, we purified hGHR-ECD (residues 1-245, omitting the signal peptide) and hGH from expression in *E. coli*. Based on CD data, the hGH was folded with the expected amount of helicity (**Suppl. Fig. 1A**). The CD spectrum of hGHR-ECD had pronounced positive ellipticities around 230 nm stemming from aromatic exciton couplings, a trait of cytokine receptors^39^, and showed as well additional contributions from disorder at 200 nm (**Suppl. Fig. 1B**). The functionality of the hGHR-ECD was confirmed from its ability to form complexes as determined from *K*_av_ for hGH, and its 1:1 and 1:2 complexes with hGHR-ECD by analytical SECs (**Fig. 1B, Suppl. Fig S1C-D**). By varying the ratio of hGH and hGHR-ECD, we could isolate the 1:2 complex and the 1:1 complex (GH in 4 times excess), and obtain the mass of hGH, hGHR-ECD and the 1:1 and 1:2 complexes using the forward scattering from SAXS and their physical extension from the derived pair-distance distribution functions, *p(r)*s (**Suppl. Table S1, Suppl. Fig. S1E-G**). Finally, to understand the ensemble properties of the hGHR-ECD in solution and generate a model, we acquired SAXS data on free hGHR-ECD at varying concentrations. The concentration normalized SAXS data overlaid perfectly (**Fig. 1C**) showing no visible interaction effects. The derived *p(r)* (insert, **Fig. 1C**) was skewed with a broad maximum around 30 Å and a maximum length (*D_max_*) of ~100 Å, consistent with the hGHR-ECD having a non-globular shape. Comparison of the SAXS data to a theoretical scattering profile obtained from one of the structures of hGHR-ECD (PDB 3HHR)^22^ resulted in a poor fit (**Fig. 1D**, blue), possibly due to the absence of the N- (1-30) and C-terminal (231-245) tails, and two disordered loops (57-61; 74-77). Next we built a model of the hGHR-ECD where the missing loops were added. The calculated scattering profile of this model provided a slightly improved fit to the SAXS data confirming that a substantial contribution to the scattering comes from disorder and conformational heterogeneity of the N- and C- terminal tails. Thus, an ensemble of 5000 models of the full-length hGHR-ECD including the N- and C-terminal tails in random configurations was built. An average of the theoretical scattering intensities from these was obtained and fitted to the experimental SAXS data (**Fig. 1D**, green). This was further refined by reweighting the ensemble against the experimental data using the Bayesian Maximum Entropy (BME) approach^41,42^, which brought χ^2^ from 34.88 to 4.67 using effectively 27% of the models, and improving the quality of the fit even further (**Fig. 1D**, red). The *R*_g_ distributions of the models before and after reweighting are shown in **Suppl. Fig. S1H**. A sub-ensemble of 500 conformations representative of the reweighted ensemble, was generated for building the model of the full-length hGHR (see below). A total of 100 conformations of this sub-ensemble is shown in **Fig. 1E**, illustrating how the disordered regions contribute considerably to the space-filling properties of the hGHR-ECD.

### The hGHR-TMD is organized parallel to the membrane normal in its monomeric state

Structures of hGHR-TMD were recently solved in dimeric states^29^ in micelles of the detergent d_38_-dodecylphosphocholine (DPC). To describe the structure and the tilt-angle of the monomeric hGHR-TMD relative to the membrane, we designed this domain of hGHR with six- and five-residues overlap with hGHR-ECD and hGHR-ICD, respectively. The resulting 36-residue hGHR-TMD (F239-R274), including an N-terminal G-S, was produced with and without isotope-labeling by a fast-track production method for single-pass TMDs^43^. Subsequently, the peptides were reconstituted in either lipid bilayers (see below) or 1,2-dihexanoyl-sn-glycero-3-phosphocholine (DHPC) micelles, successfully used for structure determination of the closely related hPRLR-TMD^30^.

A schematic overview of the extent of the hGHR-TMD α-helix determined by NMR spectroscopy and bioinformatics is shown in **Fig. 2A**. To compare the structural characteristics of this hGHR-TMD with the previously published structures^29^, we analyzed isotope-labeled hGHR-TMD in DHPC micelles by NMR and CD spectroscopy (**Fig. 2B** and **Suppl. Fig. S2A,B**). From MICS analysis^44^ of NMR backbone chemical shifts and from backbone amide *R*_2_ relaxation measurements we observed that the hGHR-TMD populated a fully formed α-helix in DHPC micelles from W249-K271 (**Fig. 2B**). This is in agreement with the findings for hGHR-TMD dimers in DPC micelles^29^, suggesting the length of the TMD α-helix to be maintained across different membrane mimetics. From the backbone chemical-shift-derived dihedral angles, a low-resolution structure to be used for building the full-length hGHR model (see below) was calculated by CYANA^45^, covering the experimentally verified helical backbone from W249-K271 (**Fig. 2C**).

**Figure 2.**
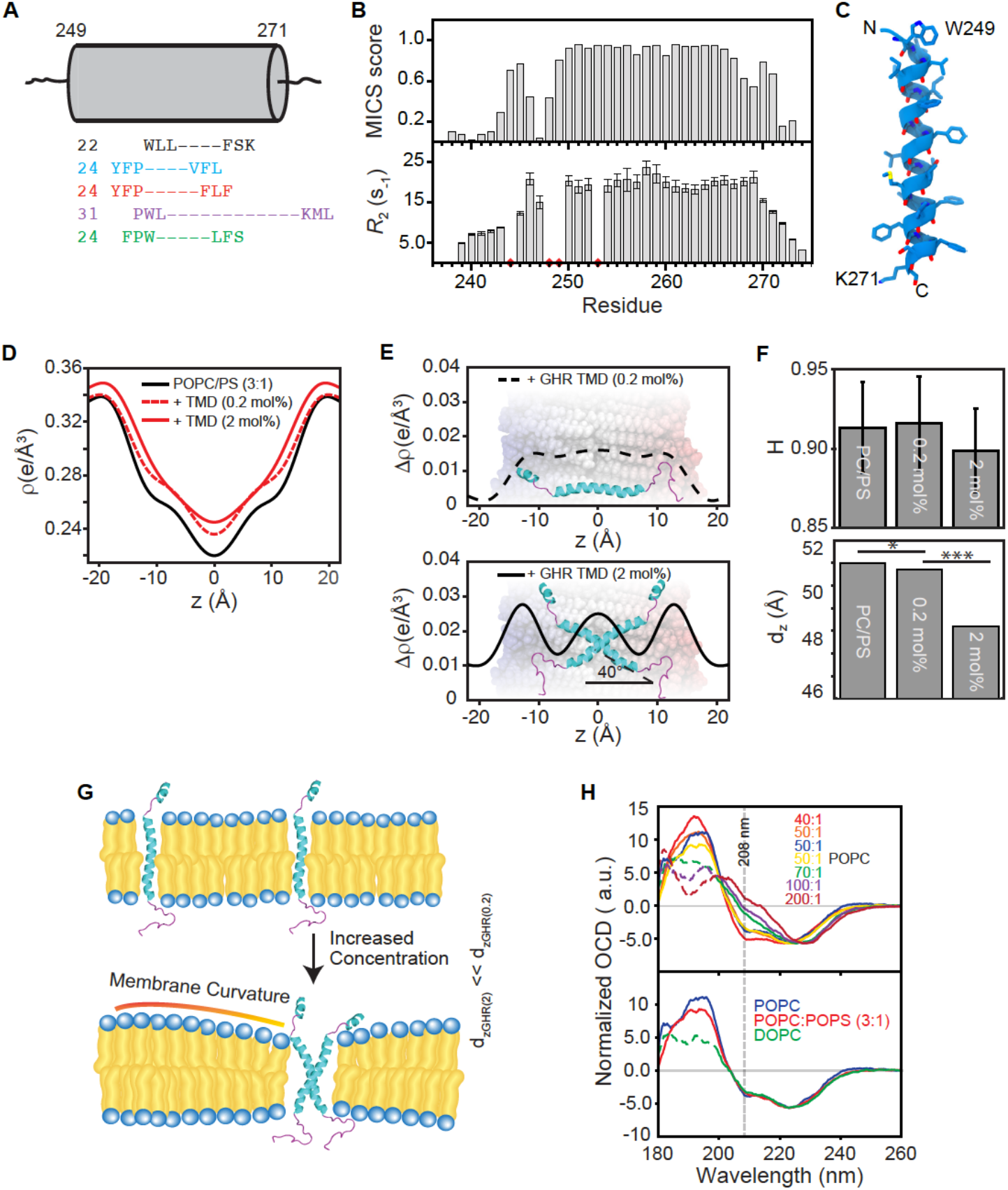
Position and definition of the single-pass α-helical TMD. (A) A schematic overview of the extent of the TMD α-helix determined by NMR spectroscopy and bioinformatic predictions. Alignment of the first three and the last residues of the TMD α-helix as determined by NMR spectroscopy (black), TMHMM^116^ (light blue), Phobius^117,118^ (red), METSAT-SVM^119^ (purple), and Uniprot annotations^120,121^ (green). The grey cylinder represents the length of the hGHR-TMD α-helix determined by NMR spectroscopy with the sequence number of the first and last residue in the α-helix. The numbers to the left of the sequences are the number of residues predicted in the TMD. (B) Top: Statistical probability for α-helical conformation as calculated by MICS^44^ based on sequence and backbone chemical shifts of hGHR-TMD in DHPC micelles, plotted against residue number. Bottom: R_2_ relaxation rates of hGHR-TMD in DHPC micelles plotted against residue number. Red diamonds highlight missing data points due to insufficient data quality or prolines. (C) Model of the hGHR-TMD α-helix. (D) Electron density profiles of lipid bilayers (POPC:POPS 3:1 mol%) with varying concentrations of hGHR-TMD (0.2 mol% and 2 mol%, respectively). (E) Difference Electron density profiles with a schematic of hGHR-TMD in an orientation best fitting to the data. (F) Illustration of membrane curvature due to monomer and dimer hGHR-TMD. (G) Top: Herman’s orientation of membranes at varying concentrations of hGHR-TMD. Bottom: Lamellar spacing of membranes at varying concentrations of hGHR-TMD. *: one-fold change and ***: three-fold change. (H) Top: OCD spectra of 6 μg hGHR-TMD in POPC, with L:P ratios varied from 1:40 to 1:200. Bottom: OCD spectra of 6 μg hGHR-TMD in POPC, POPC:POPS (3:1) or DOPC at L:P ratio 50:1. The dashed data lines represent nonreliable data due to too high HT values.

To support the modeling, we reconstituted the hGHR-TMD in a more native-like membrane system of stacked bilayers of 1-palmitoyl-2-oleoyl-sn-glycero-3-phosphocholine (POPC): 1-palmitoyl-2-oleoyl-sn-glycero-3-phospho-L-serine (POPS) (3:1 molar ratio) and investigated its structure and tilt-angle by XRD. The measured reflectivity Bragg-peaks allowed us to determine the electron density profiles, *ρ(z)*, of the different bilayer structures (**Fig. 2D**) and difference plots, *Δρ(z)*, (**Fig 2E**) of the membranes with and without inserted hGHR-TMD helices. The electron density profiles contain information about the position in the membrane and tilt angle. The electron density of the helices was calculated based on their PDB structures (PDB 5OEK; 5OHND;2N71)^29,30^ for different tilt angles and fitted to the experimental densities^46^.

The oligomeric state of single-pass TMDs was manipulated through the detergent-to-protein or lipid-to-protein (L:P) ratio^29,38^. The XRD analysis showed that at monomer conditions for the hGHR-TMD (high L:P ratio of 500:1, **Fig. 2E top**), the helix remained parallel to the membrane normal (tilt-angle 0±2°) without effects on membrane thickness, *d_z_*. At dimer conditions (low L:P ratio of 50:1) we found that the helix tilt angle changed to 40±2° relative to the membrane normal, in accordance with the GHR-dimer structures^29^ (**Fig. 2E bottom**). While the membrane flatness and intactness, as measured by Herman’s orientation function *H*, was unaffected by the presence of monomers or dimers (**Fig. 2F,** top), the dimer induced some membrane compression giving rise to a slightly thinner bilayer with smaller laminar spacing, *d_z_*, (**Fig. 2F,** bottom). An illustration of this behavior is shown in **Fig. 2G**.

To further support these observations, we employed oriented CD (OCD) spectroscopy with reconstitution of the hGHR-TMD in POPC, POPC:POPS (3:1) or 1,2-Dioleoyl-sn-glycero-3-phosphocholine (DOPC) multilamellar bilayers (**Fig. 2H** and **Suppl. Fig. S2C**). In OCD, the ellipticity of the negative band at 208 nm, which is parallel polarized to the helix axis, is strongly dependent on helix orientation, allowing distinction between a fully inserted state (I-state, parallel to membrane normal), a tilted state (T-state) or surface bound state (S-state, perpendicular to the membrane normal). At dimer conditions (L:P ratio of 50:1), the OCD spectra showed two negative bands at 208 nm and 222 nm and a positive band at 190 nm in all types of membranes tested **(Fig. 2H**), indicating successful reconstitution with formation of helical structure. Furthermore, the negative ellipticity at 208 nm was smaller compared to that at 222 nm, demonstrating the hGHR-TMD to be either in a T-state or in an equilibrium between an S-state and an I-state^47^. Increasing the L:P ratio decreased the negative band intensity at 208 nm, which even became positive at a L:P ratio of 200:1 (**Fig. 2H, top**). This indicated that at monomer conditions, the hGHR-TMD populated the more parallel I-state, fully supporting the results from XRD.

### A C-terminal GFP has no influence on the ICD ensemble

For purification of the full-length hGHR, a C-terminal GFP-H_10_-tag had to be included^48^. To ensure that this did not introduce intra- or inter-molecular interactions interfering with the hGHR-ICD ensemble, we produced the hGHR-ICD (S270-P620) without and with GFP-H_10_ (hGHR-ICD-GFP-H_10_). ^15^N-HSQC spectra of these two proteins were almost identical (**Fig. 3A**), confirming an unperturbed ensemble of the ICD. We also compared SAXS data acquired on both, which revealed a large increase in the forward scattering in the presence of GFP (**Fig. 3B**), reflecting the increase of the molar mass from 38.6 kDa for hGHR-ICD to 68.0 kDa for hGHR-ICD-GFP-H_10_ (**Suppl. Table 1**). The derived *p(r)* functions (**Fig. 3D**) showed an increased probability of short distances due to the folded GFP, but also a conserved *D_max_* consistent with an overall unaffected ICD coil conformation. The addition of GFP did not give rise to a significant change in *R_g_* (65 Å for both) (**Fig 3B**), whereas he hydrodynamic radius *(R*_h_) obtained by NMR spectroscopy, increased from 44 Å to 51 Å (**Fig 3 C**). We note that *R*_g_/*R*_h_ of ~1.5 for the hGHR-ICD falls in the range typically observed for linear chains in random coil conformations^49^ while the smaller ratio obtained for the hGHR-ICD-GFP-H_10_ is consistent with the hGHR-ICD-GFP-H_10_ containing a larger fraction of folded protein. These results taken together indicate that the C-terminal addition of GFP-H_10_ did not change the structural ensemble of hGHR-ICD.

**Figure 3.**
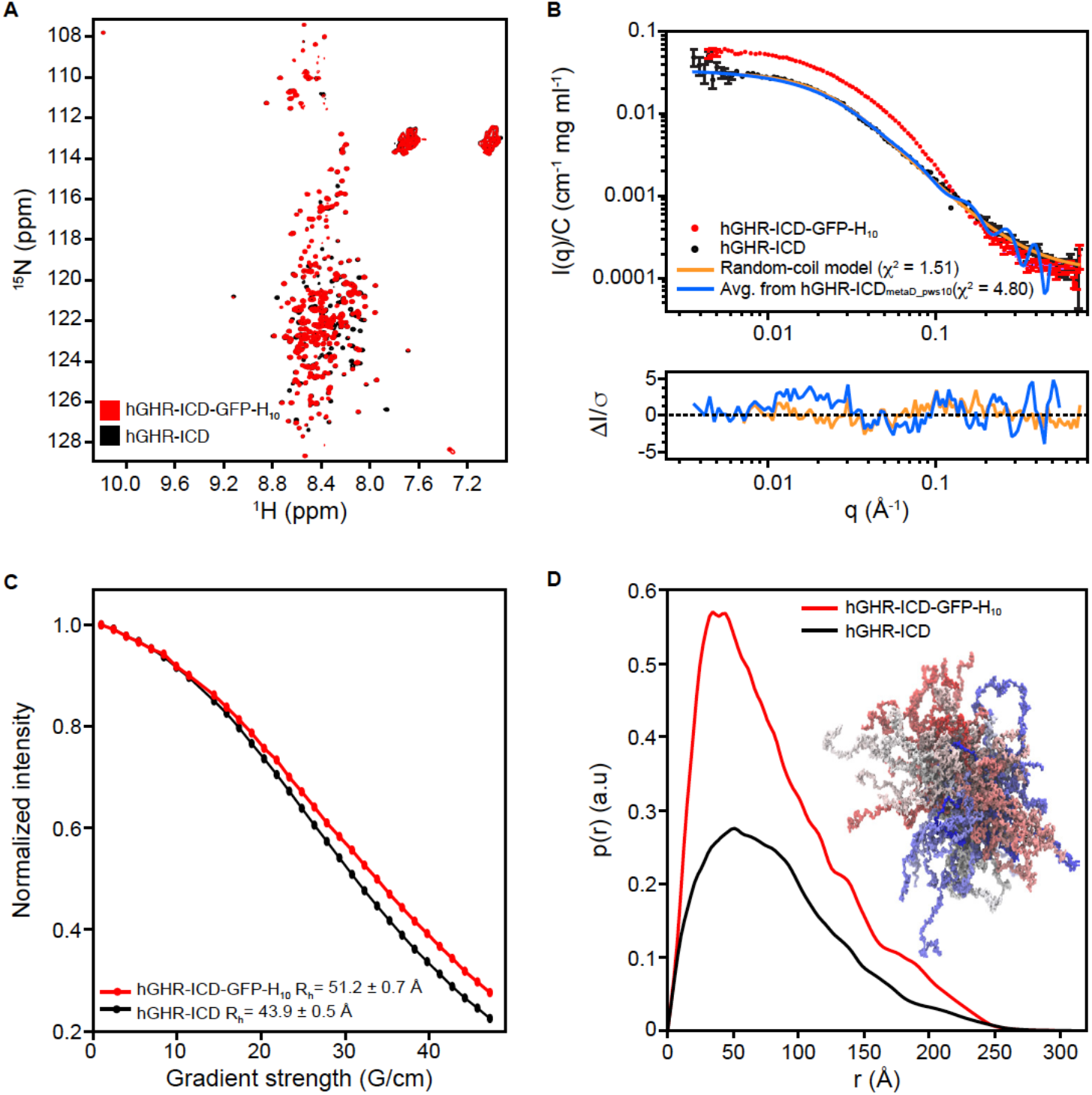
Properties of the hGHR-ICD ensemble. A) ^1^H-^15^N-HSQC spectra at 5 °C of hGHR-ICD (black) and hGHR-ICD-GFP-H_10_ (red) at 150 μM and 100 μM, respectively. (B) Concentration normalized SAXS data from hGHR-ICD (black dots, 1.1 mg/mL) and hGHR-ICD-GFP-H_10_ (red dots, 2.2 mg/mL). Fits to the data are shown for a Gaussian random coil model (orange) and from averaged scattering profiles from 1000 conformations taken from the hGHR-ICD_metaD_pws10_ simulation (1/ns) (blue). Residuals are plotted below. (C) *R*_H_ of hGHR-ICD and hGHR-ICD-GFP-H_10_ determined from pulsed-field gradient NMR. Signal decays of hGHR-ICD (black) and hGHR-ICD-GFP-H_10_ (red) are shown as a function of gradient strength together with the corresponding fits. (D) Concentration normalized *p(r)*’s derived from the above SAXS data from hGHR-ICD (black) and hGHR-ICD-GFP-H_10_ (red). An sub-ensemble of 200 conformations representative of the hGHR-ICD_metaD_pws10_ simulation is shown in the right side of the plot.

### Scaling of the protein-water interactions is required to simulate the ensemble properties of hGHR-ICD

To aid interpretation of the data of the full-length hGHR, the ensemble properties of the hGHR-ICD were modelled based on the SAXS data following two approaches: i) through fitting of the data by the form factor for simple (non-self-avoiding) Gaussian random coils^50,51^, and ii) using coarse-grained molecular dynamics simulations (CG-MD) adapted to better represent the dynamics of intrinsically disordered proteins (IDPs) to obtain an ensemble of conformations that describe the experimental data. Approach i) provided an excellent fit to the full experimental SAXS *q*-range yielding an *R_g_* of 68±4 Å (**Fig. 3B**, orange) with a χ^2^ of 1.51. This showed the average conformation of the hGHR-ICD to be very well described by a simple random coil model, which implicitly assumes a scaling exponent, *ν*=0.5. Using values empirically predicted for unfolded proteins or IDPs, or derived from computational analyses^52–55^ using slightly different scaling exponents (0.588-0.602), similar *R_g_* values of ~65Å were obtained (**Suppl. Table S2**). Hence, the values agree closely, and the effect of assuming a simple idealized Gaussian random coil model has a negligible effect on the resulting *R_g_*.

Protein-protein interactions may be overestimated in the Martini forcefield translating into unrealistic compaction of disordered regions and inability to reproduce experimentally obtained values for *R*_g_ or *R*_h_^56,57^. Recent reports investigating two three-domain protein connected by flexible linkers suggested that this could be overcome by increasing the strength of protein-water interactions^58,59^. In the case of hGHR with a long, disordered ICD, we performed unbiased and enhanced sampling MetaDynamics simulations, using the Martini 3 force field modified by increasing the strength of the protein-water interactions in the range 5-15%. Our goal was to search for a value that could provide an optimized description of the ensemble of GHR-ICD. Back-mapped atomistic conformations from these simulations were used to calculate their average R_g_ and to obtain theoretical scattering intensities, which were then fitted to the SAXS data of hGHR-ICD (**Suppl. Fig. S3**). Our results indicate that an increase in the protein-water interaction strength of 10% produced optimal results (**Fig. 3B** and **Suppl. Fig. S3**). Thus, we settled on rescaling the protein-water interaction by 10% to obtain a reliable conformational ensemble of the hGHR-ICD^1^ and to be used in the simulation of the full-length hGHR-GFP system.

### Full-length hGHR reconstituted in nanodiscs forms monomers and dimers

The intact hGHR tagged with GFP-His_10_ (hGHR) was expressed in the *S. cerevisiae* strain PAP1500, purified, and reconstituted into POPC-loaded MSP1D1 nanodiscs as described in Kassem *et al*.^48^. We used the MSP1D1 nanodisc and POPC as they are currently the most applied and best characterized carrier system by SAXS and SANS^60,61^, making computation of the nanodisc embedded full-length structure of hGHR more reliable. In SEC, the hGHR in MSP1D1 eluted over a broad peak from 10-14 mL (**Fig. 4A**). This suggested that the hGHR was reconstituted in the discs potentially as both monomers and dimers, or as higher order oligomers. To quantify the number of hGHR per disc, we performed an SDS-PAGE analysis of hGHR and MSP1D1 standards along with hGHR-loaded MSP1D1 discs isolated from the SEC at different elution volumes (**Fig. 4B**). From gel quantifications of hGHR and MSP1D1 we found that the ratio over the peak varied from ~2 hGHR per disc (F1) to ~1 hGHR per disc (F3). Since reconstitution was conducted with a 10-times excess of discs to hGHR to minimize the probability of capturing more than one hGHR pr. disc, we argue that the distribution across the peak likely represent the equilibrium between dimeric and monomeric hGHR. These results also suggested that the hGHR can form dimers in the absence of hGH as previously suggested^23^, most likely through the TMD region^23,25,62^.

**Figure 4.**
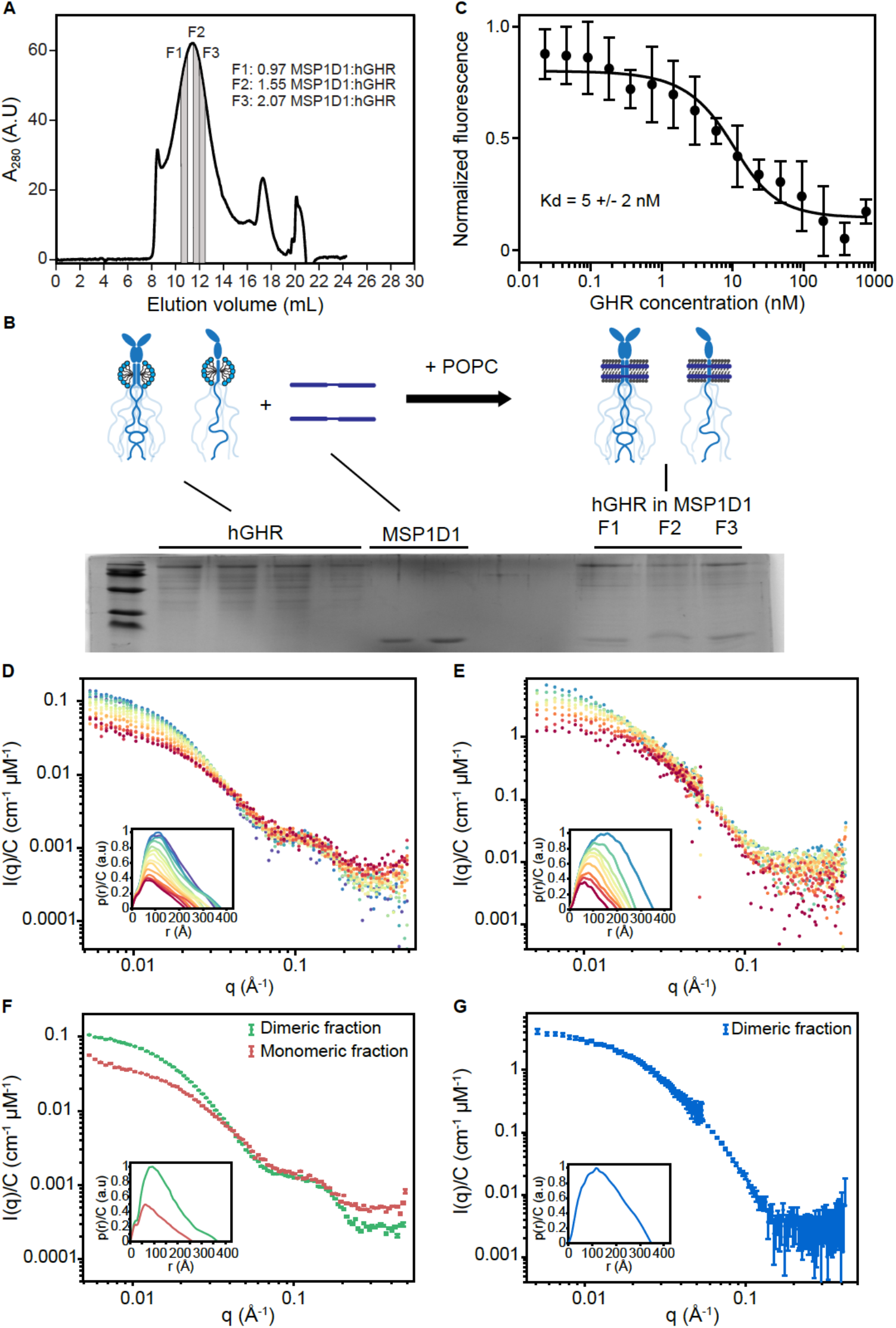
Incorporation of hGHR into MSP1D1, functional and structural analysis. (A) SEC profile of hGHR-loaded MSP1D1. The areas highlighted in grey indicate fractions (F1-F3) used for the SDS-PAGE analysis in (B). (B) SDS-PAGE analysis of hGHR and MSP1D1 standards along with hGHR-loaded MSP1D1. Fractions F1-F3 were taken from the indicated positions of the SEC purified hGHR-loaded MSP1D1 shown in (A). The illustration above the gel shows the stoichiometry of the hGHR-loaded MSP1D1. (C) MST determination of equilibrium binding constants for hGH to hGHR-loaded MSP1D1. The mean values and the standard deviation were obtained by fitting a 1:1 binding model (full line) as described in Materials and Methods. Concentration normalized (D) SAXS data and (E) SANS data of the nanodisc embedded hGHR corresponding to the highlighted SEC frames in (**Suppl. Fig. S4C,D)**. (F) Concentration normalized SAXS data from the dimer (green) and the monomer (red) fractions with the corresponding *p(r)* functions in insert. (G) Concentration normalized SANS data from the dimer fraction with the *p(r)* in insert.

### Number of lipids in the hGHR loaded MSP1D1 nanodiscs is as expected

We used phosphorus analysis^63^ performed on samples across the SEC peak (**Suppl. Fig. S4A**) to quantify the number of POPC lipids in the hGHR-nanodiscs. In the fractions with dimers (F1), the ratio between MSP1D1 nanodiscs and POPC was 115±19 and in the fraction with monomers (F3), it was 122±17. The standard deviation is based on two repetitive measurements each on two separate samples. This is comparable to results obtained in other studies of POPC nanodiscs with an α-helical membrane-anchored protein^36^ and in good agreement with the values obtained for nanodiscs solely filled with POPC (~110-130 POPC pr. nanodisc^60^). The number of lipids was used as input for the modelling of the SAXS data of hGHR-containing MSP1D1 nanodisc.

### hGHR is not N-glycosylated when produced in yeast

The hGHR has five confirmed N-glycosylation sites at N28, N97, N138, N143 and N282^64^, whereas it is unknown if it is O-glycosylated. To assess if the recombinant hGHR from *S. cerevisiae* was N-glycosylated, the electrophoretic mobility before and after treatment with endoglycosidase H was evaluated (**Suppl. Fig. S4B**). No mobility change was observed, and the band sharpness was equally high before and after treatment, suggesting lack of N-glycosylations. This is in line with previous observations on other human membrane proteins produced in the same yeast expression system^65^. To determine if yeast-produced hGHR was O-glycosylated, we performed a western blot with horse radish peroxidase conjugated with Concanavalin A that binds to mannose residues in O-glycosylated proteins^65^. A faint band corresponding to hGHR was seen indicating minor O-glycosylation (**Suppl. Fig. S4B**). As a negative control, MSP1D1 purified from *E. coli* was not detected (**Suppl. Fig. S4B**).

### Recombinant full-length hGHR reconstituted in nanodiscs is fully binding competent

To ensure that full-length hGHR embedded in the MSP1D1 nanodisc was functional, we measured equilibrium binding constants for the interaction between hGH and hGHR(MSP1D1) by microscale thermophoresis. In these studies, a 20 nM solution of fluorescently labeled (NT-647-NHS) hGH was incubated with increasing concentrations of hGHR(MSP1D1) (23 pM - 750 nM) using unlabeled hGH as control. With this approach, the dissociation constant between hGH and hGHR(MSP1D1) was determined to *K_d_* = 5±2 nM (**Fig. 4C**). As another control, we previously showed that hGHR(MSP1D1) is unable to bind human prolactin^32^, which cannot activate hGHR *in vivo*^66^. The affinities of hGH for hGHR-ECD have previously been reported as 1.2 nM and 3.5 nM for the first and the second site of hGH, respectively^67,68^. Taking all this into consideration, we find that our data agree well with previous findings and conclude that the nanodisc-reconstituted, yeast-produced full-length hGHR is fully binding competent.

### SEC-SAXS and SEC-SANS data of the full-length hGHR in nanodiscs

Structural data of the reconstituted full-length hGHR in a POPC-loaded MSP1D1 nanodisc was obtained from SEC-SAXS (**Fig. 4D**, **Suppl. Fig. S4C**) and SEC-SANS (**Fig. 4E**, **Suppl. Fig S4D**) with *p(r)* functions in inserts to **Fig 4D,E**. As was the case for the initial analysis, the SEC-elution profiles from the SEC-SAXS and SEC-SANS (**Suppl. Fig. S4C,D**) were both relatively broad and consistent with the underlying heterogeneity and systematic decrease of the particle size. Analysis of the data obtained over the SEC-SAXS and SEC-SANS elution peaks confirmed this picture, and SEC-SAXS showed a decreasing *R*_g_ from 120 Å to ~75 Å over the frames from 10-14 mL (**Suppl. Fig. 4C**). The SEC-SANS derived *R*_g_ (**Suppl. Fig. 4D**, frames from 10-14 mL) also varied over the peak, but generally less than in the SAXS experiment as a consequence of the different contrast situations in the two cases. The decrease in both the *R*_g_, the low-*q* scattering intensity and the development of the *p(r)*’s over the SEC peaks is fully consistent with the presence of discs containing first two and then one hGHR, respectively, as also supported by the initial analysis (**Fig. 4A,B**). However, in addition to dimerization, the large *R*_g_-values obtained from the left side of the SEC peak could also be affected by an overlap with the void volume (at 8-10 mL). From the data corresponding to discs with one hGHR and two hGHRs and their corresponding *p(r)*s (**Figs. 4F,G** with SEC fractions indicated in **Suppl. Fig. 4C**), a *D_max_* of ~200-250 Å was observed for monomeric hGHR in nanodiscs (SAXS only^2^). The dimeric fractions exhibited significantly larger *D_max_* of ~350 Å in both SAXS and SANS. This larger size likely results from the larger extension of the two long uncorrelated ICDs. The shoulder around 0.1 Å^−1^ of the SAXS data (**Fig. 4F)** is a typical signature of the lipid bilayer from the embedding nanodiscs^36^.

While the starting structure of a monomeric hGHR can be readily built from the chain connectivity, the structure of an hGHR dimer cannot, which complicates modeling of its structure. Further experimental complications arise both from the potential overlap with the void volume in the SEC-SAXS/SANS experiments and from possible structural heterogeneity. This may originate from a dynamic monomer-dimer equilibrium, but also from different dimers being present in the nanodisc; the biologically relevant down-down dimer conformation, a trapped up-down conformation, and even higher order structures. We therefore focused on the reliable SEC-SAXS data representing monomeric hGHR in a nanodisc and used these data to obtain the monomeric full-length hGHR structure embedded in a nanodisc bilayer.

### The structure of the monomeric full-length hGHR in a nanodisc

We followed a two-stage approach to derive a model of the structure of monomeric hGHR in the MSP1D1 nanodisc. First, we built a semi-analytical model of the nanodisc-embedded full-length hGHR (including the GFP) to refine the nanodisc parameters and validate the overall structure of the complex. Second, we used the nanodisc model from this first analysis in combination with data from a 21 μs CG-MD simulation of the hGHR embedded in a POPC bilayer. This provided an ensemble of conformations that could be back-mapped to all-atoms, and used to describe the SAXS data jointly with the refined nanodisc parameters.

The semi-analytical mathematical model of the nanodisc embedded full-length hGHR (**Fig. 5A,** see details in *Materials and Methods*) was described through four scattering amplitude components arising from, respectively, the ECD-TMD, the ICD, the attached GFP and the surrounding nanodisc. The model was implemented through the WillItFit platform^70^ and different computational approaches were applied for the different terms. In brief, the ECD-TMD, connected through a flexible linker, was represented as a rigid body through the atomic coordinates of one of the models produced with Rosetta (See *Materials and Methods*). The disordered ICD and its ensemble of conformations was modelled with a Gaussian random coil model parametrized by its *R_g_*. The attached GFP was described through its atomic coordinates (PDB 1EMA) and allowed to take a random orientation in a certain “confusion volume” in extension of the disordered ICD. For the surrounding nanodisc we allowed, as in our previous work^36,37,69^, the lipid bilayer to take a slightly elliptical shape parametrized through its axis ratio to account for the combined effect of less than maximal lipid loading and shape fluctuations. We then constrained and reparametrized the underlying geometrical model into molecular parameters such as the number of POPC per disc and the POPC area per headgroup. The scattering intensity corresponding to the model was calculated and fitted on absolute scale. An excellent model fit to the experimental data (χ^2^=1.6, **Fig. 5B**, blue) was obtained using a nanodisc containing 122 POPC lipids each with an area per headgroup of 66 Å^2^, an axis ratio of 1.5 of the elliptical bilayer and an *R_g_* of the Gaussian random coil modelling the ICD of 76 Å (see full account of model fit parameters in **Suppl. Table S3**). The number of lipids per disc was kept fixed at the value obtained from the experimental phosphorous analysis (**Suppl. Fig. S4a)**. Likewise, the axis ratio of 1.5 was fixed based on previous analyses^60^. We note that the resulting fitted POPC area per headgroup fall well within the standard disc parameters of POPC loaded MSP1D1 nanodiscs^36,69^ and that the *R_g_* of the attached ICD accords with the value we determined for the isolated ICD. The analysis shows that the semi-analytical model provides an CG low-resolution description of the nanodisc embedded GHR and form a basis for a more detailed molecular description.

**Figure 5.**
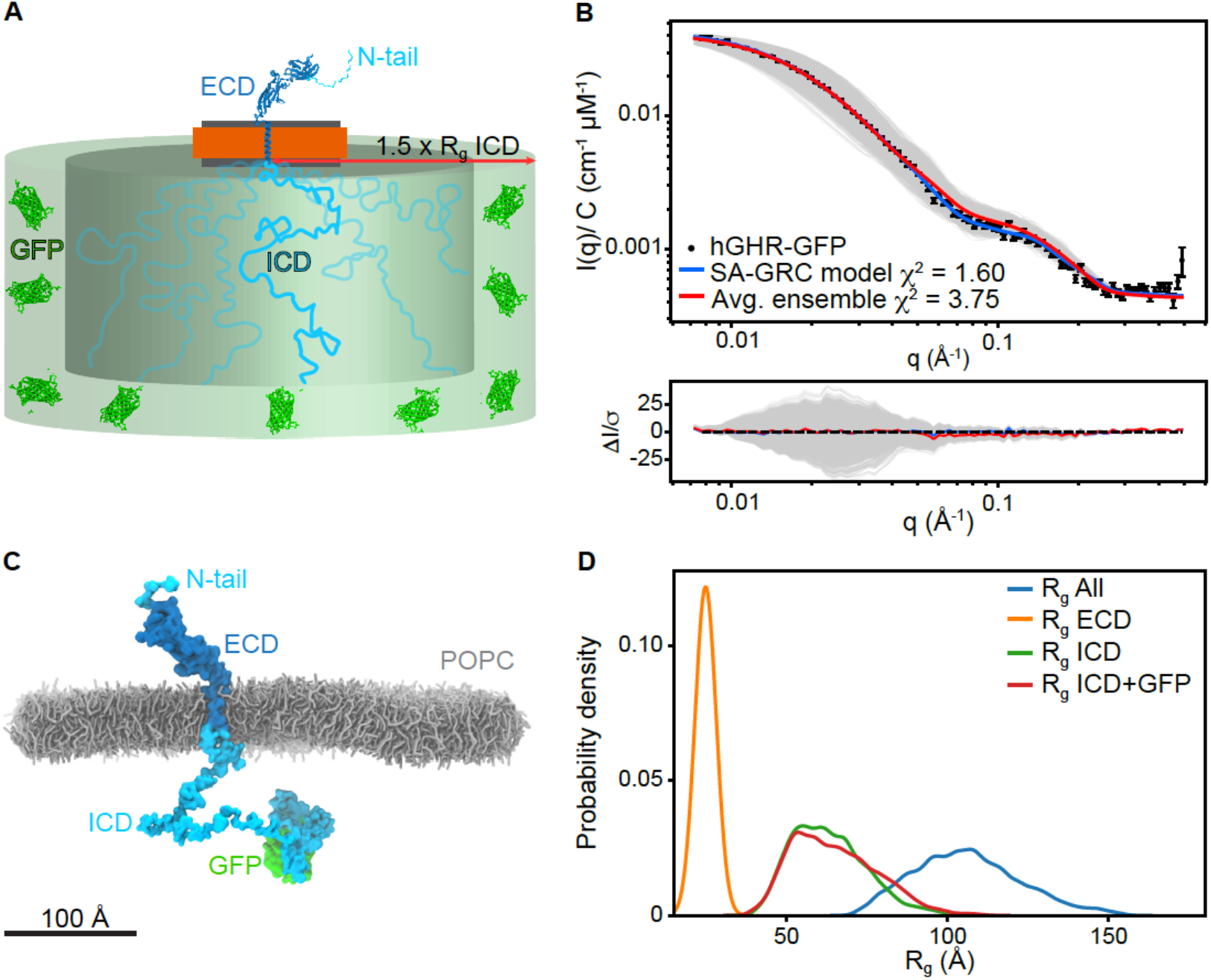
Model of the full-length hGHR in nanodiscs. (A) Schematic representation of the semi-analytical Gaussian random coil (SA-GRC) model. (B) Fits of the SA-GRC (blue) to the SAXS data of nanodisc embedded hGHR (with GFP) (blue) and of the. ensemble of 2000 conformations from the hGHR +POPC_pws10_ simulation embedded in the nanodisc (gray) and their ensemble average (red). (C) Representative snapshot from the hGHR-GFP+POPC_pws10_ simulation (*see* methods). POPC lipids shown as gray sticks, protein depicted in surface representation. Some lipids and all water and ions are omitted for clarity. (D) Probability density of the *R_g_* of the ECD (orange), ICD (green), ICD-GFP (red) and full-length protein (blue) measured from the last 20 μs of the hGHR-GFP+POPC_pws10_ simulation.

In the next stage, a CG-representation of the system was built containing the full-length receptor (residues 1-620) plus GFP (hereafter jointly named hGHR) embedded in a POPC bilayer (**Fig. 5C**). This full-length hGHR model was simulated with Martini 3 using the 10% increase in the strength of protein-water interactions found optimal for simulation of the hGHR-ICD. We simulated this system for 21 μs and extracted 2000 conformations of hGHR (one every 10 ns) from the last 20 μs. These were back-mapped to all-atom representation, and one by one embedded in the analytical nanodisc model that had been optimized through the above described semi-analytical approach and following the *WillItFit*-based procedure previously described^36,37,70^. SAXS scattering curves were calculated from the obtained ensemble (**Fig 5B**, grey), averaged (**Fig 5B**, red). We note that the average MD-derived model, despite not being refined against the experimental data in this final step, provided a very good fit to the experimental data as shown in **Fig. 5B**(red) with χ^2^ of 3.75. This confirms that the integrative approach with separate refinements of the individual domains is credible and provide a self-consistent and quantitatively correct description of the obtained data.

Further analysis of the trajectory showed that the experimental *R_g_*s obtained from the SAXS analyses of the individual parts of the protein were reproduced in the simulations which gave average *R_g_*-values of 63.3±1.2 Å for hGHR-ICD, 65.2±1.3 Å for hGHR-ICD-GFP-H_10_ and 24.9±0.04 Å for the hGHR-ECD (**Fig. 5D**). Measurement of the average helix tilt angle (15.2 ± 0.2°) (see **Suppl. Fig. S5A**), shows that the TMD remains nearly parallel to the axis normal of the membrane plane) as suggested by the XRD and OCD results obtained on the isolated hGHR-TMD. The ICD remained disordered and for the most part avoiding the membrane. Long-lived contacts and penetration of the bilayer was observed only for the intracellular juxtamembrane region (Q272-M277) and the Box1 motif (L278-K287) of the ICD, as well as for some residues from the ECD-TMD linker (**Suppl. Fig. S5B**), insert), in line with previous reports^16^. Visual inspection of the trajectory (**Suppl. movie M1**) showed that the ECD-TMD linker remained flexible allowing the ECD to adopt a range of orientations while remaining mostly upright as shown by the angle between the principal axis of the D2 domain and the z-axis (average 36.8 ± 0.7°, **Suppl. Fig. S5C**). We note that the D1 domain remained far from the lipid surface. The N-terminal tail of the ECD remained disordered without long-lived contacts with the folded part of the ECD or the membrane.

In summary, the integrative model of the full-length monomeric hGHR in a nanodisc, containing almost equal amounts of structural order and disorder, fully captured the SAXS data recorded on the complex molecular system. Hence, the model provides the first molecular insight into the structure of an intact, full-length class 1 cytokine receptor in a lipid membrane carrier system.

## DISCUSSION

Membrane proteins take on a variety of different topologies, sizes and functions and a large portions of membrane proteins exist in tripartite structures that require different handling schemes and methodological studies. Such complexities are further amplified for membrane proteins having large fractions of structural disorder^32,71,72^, which impose obstacles to classical structural biology. Thus, different topologies and order/disorder dispositions require different approaches, and one particular group of membrane proteins falls between the cracks by being too small and unstructured for cryo-EM, too large for NMR spectroscopy and too dynamic for X-ray crystallography. An important subgroup of these membrane proteins, which plays key biological roles, is the cytokine receptor family.

In the present work, we examined the structure of an archetypal and particularly challenging membrane protein, the cytokine receptor hGHR, for which 50% of its chain is intrinsically disordered (**Fig. 6**). The structure of the monomeric hGHR revealed that when inserted in a bilayer mimetic, neither the ECD nor the long, disordered ICD engage in long-lived contacts with the membrane. This is remarkable, although it should be noted that the lipids used in the current study are not fully mimicking the complexity of native membranes lacking phosphoinositides or/and cholesterol, just as the proteoglycan layer on the extracellular side and the cytoskeleton on the inside is missing. We did, however, capture some lipid interactions by the intracellular juxtamembrane region (**Suppl. Fig. S5B**), which have been previously described^16^. It is possible that the native composition of the bilayer may influence the conformation of the receptor, but inherently there is no affinity for the POPC bilayer. Thus, the intracellular, disordered domain protrudes away from the bilayer and into the cytosol. Its average *R_g_* of 65-70 Å corresponds to an average end-to-end distance of about twice this value. This defines its capture distance and the large search volume (**Fig. 6** and **Suppl. Fig. S5D**), which allows it to scout for and engage with kinases, phosphatases and regulatory proteins such as the signal transducer and activator of transcription (STAT)s, suppressors of cytokine signaling (SOCS)s and the cytoskeleton^14^.

**Figure 6.**
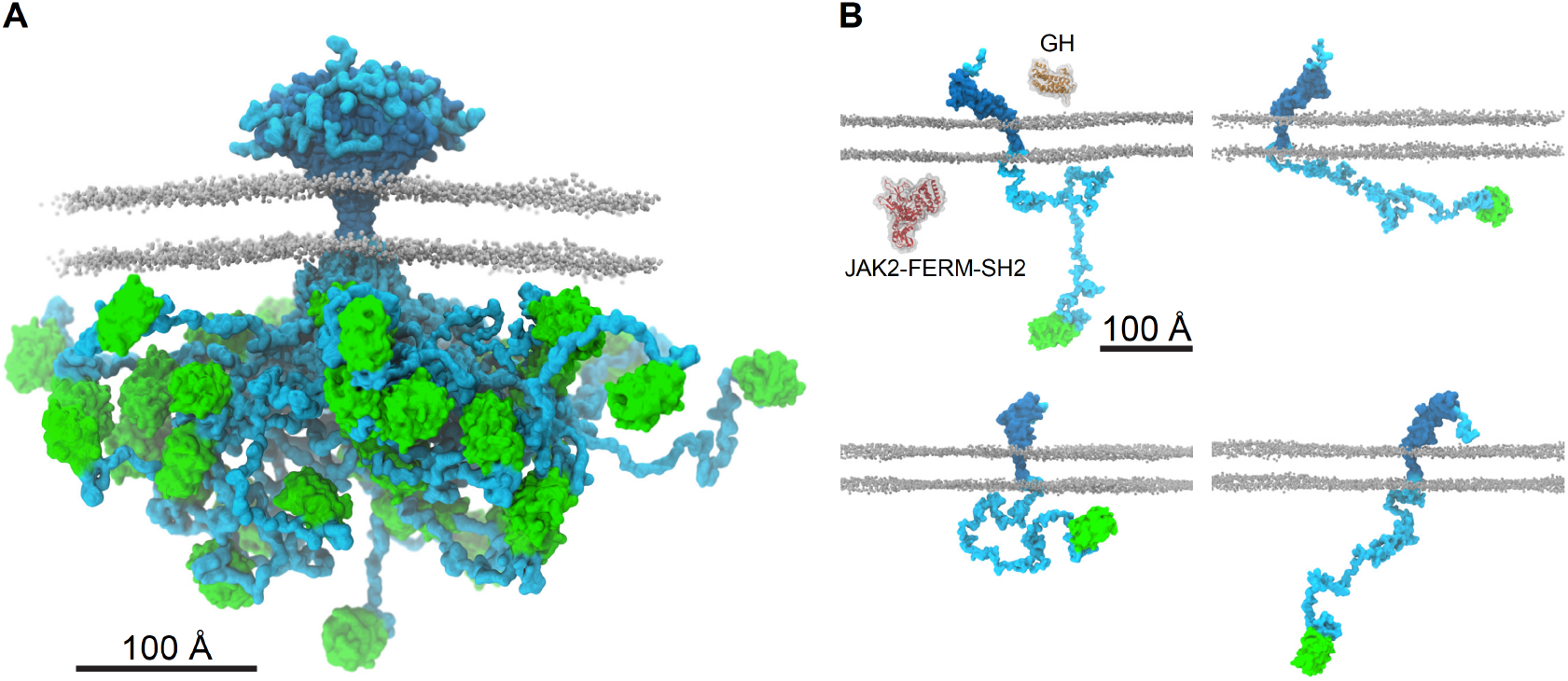
The ensemble structure of membrane embedded full-length human GHR. (A) Representative ensemble of conformations obtained from the last 20 μs of the hGHR-GFP+POPC_pws10_ simulation. Color scheme and representations as in Figure 5C. (B) Examples of the multitude of domain orientations of hGHR in the membrane. In the first panel, the structures of hGH (PDB 3HHR_A, orange) and of JAK2-FERM-SH2 (PDB 4Z32, red) are shown. Color scheme and representation of hGHR and POPC as in Figure 5C.

A particular noteworthy observation from the structure of hGHR is the disordered, ~30 residue N-terminal of the ECD, which has been neglected in all previous structural studies. The role of this N-terminal IDR in GHR function is unknown, but N-terminal IDRs are present in other family members, including the EPOR. An isoform of the GHR with a 22 residues deletion in the disordered N-tail (*d3*-GHR) shows altered ERK1/2 signaling but unaltered STAT5 signaling, and *d3*-GHR individuals show an increased lifespan^73^. Thus, key functional relevance is coupled to the N-tail. A search in the eukaryotic linear motifs (ELM) database^74^ suggests the presence of a glycosaminoglycan attachment site, _1_FGFS_4_, in the tail. Of relevance to this, the WSXWS motif, which in hGHR is YGEFS, constitutes a C-mannosylation site linking the C1 atom of the α-mannose to the indole C2 atom of the tryptophan^75,76^. The WSXWS motifs has also been suggested to bind GAGs^21^, so it is possible that the disordered N-tail of hGHR play similar roles as the WSXWS motif, and we notice a degenerate motif of this kind, also in the N-tail, given by the sequence _16_WSLQS_20_. Nonetheless, the function of the disordered N-tail of hGHR remains unestablished.

The integrative nature of our approach to determine the structure of the hGHR required development and optimization of several protocols. This was particularly necessary during the modelling and fitting of the SAXS data based on the combined semi-analytical and experimentally driven molecular modelling approach to account for the structure and large flexibility of the hGHR. Key to the success was a scaling of the strength of the protein-water interaction in the CG molecular dynamics simulations of the ICD and full-length hGHR. This enabled reliable fits to the disordered chain in terms of *R_g_*. On the semi-analytical modelling side, we have expanded our previous approaches to interpret scattering data from bare nanodiscs and rigid membrane proteins incorporated into these^36,37,69^, to now also allow for modelling membrane proteins with significant amounts of intrinsic structural disorder. We emphasize that even if the parameters of the GHR model are custom fitted to the hGHR system, the approach is fully generalizable and may be adapted to membrane proteins of similar topology provided that high quality SAS data are available. Thus, the use of this integrative semi-analytical and MD simulation-based approach suggests that SAS in combination with MD simulations is a useful way of retrieving structural models to provide structural insight into otherwise “method orphan” membrane proteins, in particular highlighting the interdomain orientations. This opens the door for more systematic investigations of for example single-pass transmembrane proteins in different environments, e.g. with respect to the lipid composition, the buffer environment or with binding partners to understand how these very dynamic membrane proteins transduce information across the membrane. This additionally includes other single-pass membrane proteins with similar complexity such as the cadherins and cell adhesion molecules (e.g. downs syndrome cell adhesion molecule), but also membrane proteins with long disordered regions such as the solute carrier family 9, type II receptor serine/threonine family, and palmitoyl transferases^48,77^.

A key observation made possible from acquiring data on the full-length hGHR, is the lack of restriction on the relative orientation of the domains (**Fig. 6**). Not only is the ICD and the N-tail disordered, but the flexible linker joining the ECD and TMD combined with the lack of membrane association allow them to freely reorient relative to each other, at least in the free state (**Fig. 6**). Thus, in addition to structure, it becomes important to consider how the flexibility of the entire chain take on roles in signaling. From our studies we were not able to derive if correlated motions between the ECD and the ICD exist. However, once the hGH binds to the ECD, changes in conformation and flexibility may propagate along the chain reaching the ICD and bound protein partners, eliciting signaling. Similar suggestions were put forward based on data from solid-state NMR studies on the epidermal growth factor receptor, revealing increased dynamics in the bound state^78^. Since the JAK2 binding site only constitutes ~6% of the ICD, and the STAT5 docking sites are ~200-300 residues away from it^79^, conformational changes involving redistribution of the structural ensemble of the long, disordered region need to be achieved in a controlled manner. It is currently unclear how this is accomplished, but phosphorylations or binding to other proteins are likely to impact the ensemble, including the degree of compaction. Finally, the long ICD has a high content of short linear motifs (SLiMs), which are distributed along the chain in SLiM hotspots^15^, and the space occupied by the free ICD (**Suppl. Fig. S5E,F**) may therefore enable room for generation of larger, supramolecular signaling complexes constituted by many partners. So far, only binary complexes involving the ICD have been considered. With the presence of two disordered chains in a dimer, the occupied space of each ICD chain is reduced due to steric exclusion, which may result in different supra-molecular complexes compared to those involving the monomer. Thus, understanding the role of structural disorder in orchestrating cellular signaling by disorder remains enigmatic. With the first structure of a full-length membrane protein embedded in a realistic membrane scaffold and containing a large disordered chain at hand, the understanding of regulation of signaling by disordered chains, often present in higher order assemblies of several chains, now has a molecular platform from which new questions can be tackled.

## Acknowledgments

This work has been supported by the Novo Nordisk Foundation Synergy programme (BBK, LA), and Challenge Program (REPIN, BBK), the Lundbeck Foundation (BBK) and the Lundbeck Foundation Initiative BRAINSTRUC (KLL, BBK, LA). The authors acknowledge the ILL, France, for the allocated SEC-SANS beam time as well as the European Synchrotron Radiation Facility (ESRF) and PETRAIII at the Deutsches Elektronen-Synchrotron (DESY), Germany for the allocated SAXS beamtime. Furthermore, we thank Signe A. Sjørup and Jacob Hertz Martinsen for skilled technical assistance, Nicolai Tidemand Johansen for help in relation to nanodisc preparations, SAXS and SANS measurements and data reduction. We also thank the D22 technicians Mark Jacques, Anne Martel, Lionel Porcar, the beamline scientists Marta Brennich and Petra Pernot at BM29 and beamline scientist Hayden Mertens at P12 for technical support during beamtimes, including pilot beamtimes.

## Data availability

All data generated and analyzed in this study will be made available as source data upon publication of the manuscript. SAS data will be uploaded to the SASBDB data base. Representative subsections of the MD data will be made available on github (https://github.com/Niels-Bohr-Institute-XNS-StructBiophys) while the full sets will be made available upon request to the authors.

## Code availability

All codes utilized in this study are available from the authors upon request. The implemented *WillItFit* routines are open source and will be made accessible as an update on the *WillItFit* repository at Sourceforge: https://sourceforge.net/projects/willitfit/

## Author contribution

N.K., R.A-S., K.B., M.R., P.A.P., L.A., B.B.K. designed the research. N.K., R.A-S., K.B., A.B., H.S., A.K., A.J.L., J.B., and A.S.U. performed research and/or contributed new reagents. N.K., R.A-S., K.B., A.B., H.S., A.K., J.B., M.C.P., Y.W., M.C.R., P.A.P., K.L-L., L.A., and B.B.K. analyzed data. N.K., R.A-S, K.B., L.A., and B.B.K. wrote the paper with input from all authors.

## Competing interests

The authors declare no competing interests.

## MATERIALS AND METHODS

### hGHR-ECD expression and purification

The DNA sequence coding for hGHR-ECD (1-245, C242S, no signal peptide) in a pET11a was bought from Genscript and transformed into competent Rosetta2 (DE3)pLysS cells. These were grown in 1 L LB medium with 3 % (v/v) ethanol, containing 100 ug/mL ampicillin and chloramphenicol to OD_600_ = 0.6-0.8, and induced by addition of 0.5 mM of IPTG for 4 h at 37°C and 160 RPM. The cells were harvested by centrifugation (5000 x g for 15 min) and resuspended in 1x PBS (140 mM NaCl, 2.7 mM KCl, 10 mM Na_2_H_2_PO_4_, 1.8 mM KH_2_PO_4_), pH 7.4 containing 25 % (w/v) sucrose and 5 mM EDTA. The cells were lysed on ice by sonication using an UP400S ultrasonic Processor, 6×30s sonication followed by 30s rest at 50% amplitude. Following centrifugation (20,000 ×g, 4°C) for 25 min, the pellet was resuspended in 1x PBS pH 7.4, containing 25 % (w/v) sucrose and 5 mM EDTA, repeated three times in total. The pellet was solubilized in 500 mL 50 mM Tris-HCL pH 8.5, 10 mM beta-mercaptoethanol (bME), 6 M urea and heated for 5 min at 55 °C and left O/N with slow stirring, 4°C. The amount of hGHR-ECD was estimated on an SDS PAGE by comparing to the LMW and diluted to a concentration below 0.1 mg/mL in 50 mM Tris-HCL pH 8.5, 10 mM bME, 6 M urea. To refold, hGHR-ECD was dialyzed against 4 L 150 mM NaCl, 50 mM Tris-HCL pH, 8.5, 10/1 mM cysteamine/cystamin at 4°C, 12 kDa MW cut off until the urea concentration was below 0.1 M. Following centrifugation at 20,000 ×g for 15 min, the sample was placed on ice and stirred slowly while ammonium sulphate was added to a final concentration of 75 % (w/v) and then left for two hrs. The solution was centrifuged at 12,000 ×g at 4°C, 25 min, and the pellet dissolved in 100 mL miliQ water and left for 2 h, followed by dialysis against 30 mM NH_4_HCO_3_, pH 8.0 overnight at 4°C. After centrifugation at 13,000 ×g for 15 min, the supernatant was concentrated using a Millipore spinfilter (10 kDa cut-off), and applied to a Superdex 75 16/85 column (GE health care) at 4°C, 150 mM NaCl, 30 mM NH_4_HCO_3_, pH 8.5. Selected fractions were reapplied to a Superdex 200 increase 10/300 GL column in 20 mM Na_2_H_2_PO_4_, pH 7.5, 150 mM NaCl, prior to SAXS measurements.

### hGHR-ICD expression and purification

The coding region for hGHR-ICD (S270-P620) was cloned into a pGEX-4T-1 vector, containing an N-terminal GST-tag followed by thrombin cleavage site and transformed into Bl21(DE3) cells. Expression was done in 1L Terrific Broth (TB) medium containing 100 ug/mL ampicillin. At OD_600_ = 0.6-0.8 cells were induced by 1 mM of IPTG for 4 h, at 37°C and 160 RPM. Cells were harvested by centrifugation and resuspended in 40 mL 1x PBS, pH 7.4, 0.1 % (v/v) Triton X-100 and a tablet complete EDTA-free protease inhibitor cocktail. The cells were lysed on ice by sonication using an UP400S ultrasonic Processor, 4 times 30s sonication followed by 30s rest at 100% amplitude. Following centrifugation (20,000 ×g, 4°C) to remove cellular debris, the lysate was applied to a Glutathione Sepharose 4 Fast Flow column (GE health care) and incubated for 2 h at 25 °C. The column was washed with 50 mL 1x PBS, pH 7.4 and eluted 20 ml 50 mM Tris-HCl, 10 mM reduced glutathione, pH 7.4. The eluted solution was dialyzed against 1 L 20 mM Tris-HCl, 150 mM NaCl, pH 7.4 at 4 °C. The GST-tag was cleaved off by the addition of 20U thrombin /L culture, leaving residues GS in the N-terminal. The sample was then concentrated, 10 mM DTT added and heated to 72°C for 5 min, incubated on ice, and centrifuged for 20,000 ×g at 4°C for 10 min. A final purification on a Superdex 200 increase 10/300 GL column (GE Healthcare) in 20 mM Na_2_H_2_PO_4_, pH 7.5, 150 mM NaCl was done and selected fractions were used for SAXS measurements.

### hGHR-ICD-GFP-H_10_ expression and purification

The coding region for hGHR-ICD (S270-P620) including an N-terminal methionine, C-terminal TEV cleavage (ENLYFQS) site followed by a yeast enhanced GFP^80^ and 10 histidines (hGHR-ICD-GFP-H_10_) in a pET-11a vector was bought from GeneScript. Expression was done in 1L Terrific Broth (TB) medium (for SAXS) and in ^15^N-labeled minimal medium (22 mM KH_2_PO_4_, 62.5 mM NaH_2_PO_4_, 85.6 mM NaCl, 1 mM MgSO_4_, 1 ml “trace element solution”, 4 g glucose, 1.5 g NH_4_Cl (^15^N labelled nitrogen)) (for NMR) containing 100 ug/mL ampicillin. At OD_600_ = 0.6-0.8, expression was induced by 1 mM of IPTG for 3 h, at 37°C and 160 RPM. Cells were harvested by centrifugation and resuspended in 40 mL 1x PBS, pH 7.4, and a tablet complete EDTA-free protease inhibitor cocktail. The cells were lysed on ice by sonication using an UP400S ultrasonic Processor, 4 times 30s sonication followed by 30s rest at 100% amplitude. Following centrifugation (20,000 ×g, 4°C), the pellet containing hGHR-ICD-GFP-H_10_ was solubilized by adding 40 mL 20 mM NaHCO_3_ pH 8.0, 150 mM NaCl and 8 M urea. Following centrifugation (20,000 ×g, 4°C), the supernatant was refolded by dialysis in two steps. First by dialysis in 4 L 20 mM NaHCO_3_ pH 8.0, 150 mM NaCl, 4 M urea at 4°C using 3 kDa molecular weight dialysis bag cut-off for 4 h, and then in 4 L 20 mM NaHCO_3_ pH 8.0, 150 mM NaCl at 4°C overnight. Following centrifugation (20,000 ×g, 4°C), the supernatant was applied to a prepacked 5 mL Ni-resin column. The column was washed with 3 column volumes (CV) of 20 mM NaCHO_3_ pH 8, 150 mM NaCl, 10 mM imidazole and eluted using 20 mM NaCHO_3_, pH 8.0, 150 mM NaCl, 250 mM imidazole. Fractions containing hGHR-ICD-GFP-H_10_ were concentrated and applied to a Superdex 200 16/60 increase column in 20 mM NaH_2_PO_4_/Na_2_H_2_PO_4_, pH 7.5, 150 mM NaCl. Fractions containing hGHR-ICD-GFP-H_10_ were analysed by SDS-PAGE and selected fractions were used for SAXS and NMR experiments.

### hGH purification

hGH in a pJExpress414 was bought from ATUM, USA (formerly known as DNA2.0) and transformed into competent BL21 (DE3) cells. These were grown in 1L TB containing 100 ug/mL ampicillin to OD_600_ = 0.6-0.8 and induced by addition of 1 mM of IPTG for 4 h, at 37°C and 160 RPM. Cells were harvested by centrifugation (5,000 xg, at 4°C, 25 min) and resuspended in 50 mL 50 mM Tris, 0,5 mM EDTA, pH 8.0, 1 mM PMSF. Cells were lysed on ice by sonication using an UP400S ultrasonic Processor, 5 times 30s sonication followed by 30s rest at 50% amplitude. Following centrifugation at 10,000 ×g, at 4°C, 15 min, the pellet was resuspended in 20 mL 10 mM Tris, 1 mM EDTA, pH 8.0, 1 mM mM PMSF. The pellet was re-centrifuged two times, the supernatant discarded, and solubilized in 250 mL 5 M guanidinium chloride (GuHCl), 200 mM Na_2_HPO_4_/NaH_2_PO_4_, pH 7.0, 15 mM bME. The solution was heated for 10 min at 55 °C and stirred mildly for 2 h at room temperature. The solution was diluted in denaturation buffer (5 M GuHCl, 200 mM Na_2_HPO_4_/NaH_2_PO_4_, pH 7.0, 15 mM bME to reach a hGH protein concentration below 0.1 mg/mL. The solution dialysed in a 5 L beaker, with a drain in the top, filled with 5M GuHCl, 200 mM Na_2_HPO_4_/NaH_2_PO_4_, pH 7.0, 15 mM bME. A peristaltic pump was used to add refolding buffer (20 mM NH_4_HCO_3_ pH 8.0, 200 mM NaCl) in the bottom of the beaker with a flowrate of 1.5 mL/min. After three days, when the GuHCl concentration was below 1.5 M, the dialysis bags were transferred to a new 5 L beaker with 20 mM NH_4_HCO_3_ pH 8.0, 200 mM NaCl, and dialysed three times until the concentration of GuHCl was below 0.1 M. Following centrifugation for 18000 ×g for 10 min, the supernatant was concentrated using a Millipore Pellicon module to approximately 30 mL. The solution was applied to a Superdex75 26/600 column in 20 mM NH_4_HCO_3_, 100 mM NaCl, pH 8.0. Selected fractions were dialysed against 5 L 20 mM Tris pH 8.0 twice, and applied to a HiTrap QFF 5mL. The sample was eluted in 20 mM Tris pH 8.0 by a salt gradient from 0-1M NaCl at a flow rate of 5 mL/min over 20 CV. Selected fractions were flash frozen in liquid nitrogen and left at −20 prior to use.

### Analytical SEC

Analytical SEC experiments of a set of samples with various ratios of hGH:hGHR-ECD were run on Superdex 200 increase 10/300 GL column in 20 mM Na_2_HPO_4_/NaH_2_PO_4_ pH 7.4, 100 mM NaCl at room temperature with a flowrate of 0.5 mL/min. Protein sample concentration were in the micromolar range but varied. The column was calibrated using conalbumin (75 kDa), ovalbumin (44 kDa), carbonic anhydrase (29 kDa), ribonuclease A (13.7 kDa), acetone and blue dextran and apparent partition coefficient, ***K***_AV_, was determined for all peaks.

### Circular dichroism spectroscopy

Far-UV CD spectra were recorded on 10 μM hGHR-TMD in 2 mM DHPC, 5 μM hGH and 5 μM hGHR-ECD in 10 mM Na_2_HPO_4_/NaH_2_PO_4_ (pH 7.4). The spectra were recorded on a Jasco J-810 Spectropolarimeter in a 1 mm quartz glass Suprasil cuvette (Hellma) at 20°C. A total of 10 scans were accumulated from 260 nm to 190 nm for each sample and buffer background was recorded at identical setting and subtracted. For hGHR-TMD, the background included 2 mM DHPC. The scan mode was continuous with a speed of 10 nm/min and a data pitch of 0.1 nm. The spectra were processed and smoothened (means-movement method, convolution width 25) and converted into mean residue ellipticity values.

### hGHR-TMD purification

hGHR-TMD was expressed in *E. coli* and purified as previously described^43^.

### Oriented circular dichroism

hGHR-TMD was dried under a flow of N_2_ and subsequently dissolved in MeOH:CHCl_3_ (5:1) to reach a final stock solution of 0.4 mg/ml hGHR-TMD. To validate the concentration, 100 μl of the stock solution was dried under N_2_ flow and resuspended in 100 μl 50 mM SDS in PB buffer (pH 7.0) and the absorbance at 280 nm was measured. Lipid stock solutions of POPC, DOPC and POPC/POPS (3:1) were prepared in MeOH:CHCl_3_ (1:1) at 0.25 mg/ml and 5 mg/ml. The protein and lipid stock solutions were mixed in the following L:P ratios; 40:1, 50:1, 70:1, 100:1, 150:1 and 200:1. 6 μg protein was applied to a quartz glass with a Hamilton pipette for each experiment. The sample was spread over a fixed circular area on the glass and subsequently dried under vacuum for 3 h to remove the MeOH:CHCl_3_. The dried sample was mounted in a sample holder and was hydrated overnight in a chamber with a saturated K_2_SO_4_ solution at 20 °C. Finally, the samples were loaded into a rotor in a Jasco J-810 spectropolarimeter and spectra were recorded from 8 different angles; 0, 45, 90, 135, 180, 225, 270 and 315°. Each spectrum was measured twice from 260 to 180 nm with a scanning speed of 20 nm/min, a data pitch of 0.1 and a response time of 8 s. The spectra were averaged and reference OCD spectra from samples with the same amount of lipid was subtracted. The OCD spectra were recorded from 8 different angles to even out linear dichroism^47^ (Suppl. Fig. S2C). The spectra from different angles were averaged and background-subtracted and normalized to the intensity at 222 nm. High voltage effects prevented the measurement of higher L:P ratios.

### X-Ray Diffraction

Highly-oriented multi lamellar membranes were prepared on single-side polished silicon wafers. POPC (Avanti), POPS (Avanti), and 1,2-dimyristoyl-sn-glycero-3-phospho-L-serine (DMPS, Sigma) were mixed with hGHR-TMD at 2 and 20 mol% concentrations in 2,2,2-trifluoroethanol:chloroform (1:1, vol/vol) at a solution concentration of 18 mg/mL. The wafers were sonicated in 1,2-dichloromethane for 30 min, and then rinsed with alternating methanol and 18.2 MΩ ⋅ cm water. The wafers were dried, and 75 μL of solution was deposited. After drying, the samples were placed in a vacuum for 24 h at 37 °C to allow for trace solvent evaporation and annealing. Samples were then hydrated in a closed chamber at 97% RH with a separate K_2_SO_4_ saturated solution for 48 h prior to scanning.

XRD data was obtained using the Biological Large Angle Diffraction Experiment (BLADE) at McMaster University. BLADE uses a 9 kW (45 kV, 200 mA) CuKα rotating anode at a wavelength of 1.5418 Å using a Rigaku HyPix-3000 2D semiconductor detector with an area of 3000 mm^2^ and 100 μm pixel size^81^. All samples were prepared and measured in replicates to check for consistency. Electron density profiles were determined from specular reflectivity, as previously described^46^. The lamellar spacing, *d_z_*, was determined from the spacing of the reflectivity Bragg peaks. Herman’s orientation function was determined by integrating the intensity of the 3^rd^ Bragg peak as function of the meridonal angle *ϕ* (the angle relative to the *q_z_* axis), as described in^82^.

### NMR spectroscopy

NMR spectra were recorded on a 750 MHz (^1^H) Bruker AVANCE spectrometer equipped with a cryogenic probe. Unless otherwise specified, all NMR samples contained 10 % (v/v) D_2_O and 1 mM 4,4-dimethyl-4-silapentane-1-sulfonic acid (DSS). Proton chemical shifts were referenced internally to DSS at 0.00 p.p.m., with heteronuclei referenced by relative gyromagnetic ratios. Free induction decays were transformed and visualized in NMRPipe^83^ or Topspin (Bruker Biospin) and analysed using CcpNmr Analysis software^84^. For hGHR-TMD, all NMR spectra were recorded at 37 °C in 2 mM tris(2-carboxyethyl)phosphine (TCEP), 0.05% (v/v) NaN_3,_ 50 mM NaCl and 20 mM Na_2_HPO_4_/NaH_2_PO_4_ (pH 7.4). The spectra for backbone assignments of hGHR-TMD (HNCO, HNCAHC, HNCA, HNCACB, CBCA(CO)NH, ^1^H, ^15^N-HSQC) were measured on 1 mM ^13^C,^15^N-hGHR-TMD solubilized in 210 mM DHPC. Secondary structure content was evaluated from backbone chemical shifts using the motif identification from chemical shifts (MICS) programme^44^. *R*_2_ transverse relaxation rates of 0.5 mM ^15^N-hGHR-TMD in 110 mM DHPC were determined from a series of ^1^H,^15^N-HSQC spectra with varying relaxation delays between 10 and 250 ms and triple replica at 130 ms. The relaxation decays were fitted to single exponentials and relaxation times determined using CcpNmr Analysis software^84^. A low-resolution model of hGHR-TMD was calculated using CYANA^45^ including only dihedral angles restraints from TALOS^85^ using the backbone chemical shifts. Standard settings were used calculating 50 conformers with 4000 torsion angle dynamics steps. The 10 best conformers, with the lowest CYANA target function score was used for further modelling.

All NMR data of hGHR-ICD and hGHR-ICD-GFP-H_10_ were acquired at 5°C to minimize amide exchange in 1 mM TCEP, 150 mM NaCl and 20 mM Na_2_HPO_4_/NaH_2_PO_4_ (pH 7.4). ^1^H,^15^N-HSQC spectra were acquired at concentrations of 150 μM for ^15^N-hGHR-ICD and 100 μM for ^15^N-hGHR-ICD-GFP-H_10_. The hydrodynamic radii (*R_H_*) of hGHR-ICD and hGHR-ICD-GFP-H_10_ were determined from a series of ^1^H, ^15^N-HSQC spectra with preceding pulse-field gradient stimulated-echo longitudinal encode–decode diffusion filter^86^ and with the gradient strength increasing linearly from 0.963 to 47.2 G cm^−1^. To determine the diffusion coefficients (*D*) the decay curves of the amide peaks were plotted against the gradient strength and fitted in Dynamics Center (Bruker) using *I* = *I_0_*exp(–D_x_ ^2^*γ*^2^*δ*^2^(*Δ* − *δ*/3) ×10^4^), in which *I* is the intensity of the NMR signal at the respective gradient strength, *I_0_* the intensity without applied gradient, x the gradient strength in G cm^−1,^ *γ* = 26752 rad Gs−1, *δ* = 3 ms, *Δ* = 250 ms. *R_H_* was calculated from the diffusion coefficient using the Stokes–Einstein relation, *R_H_* = k_B_T/(6π*ηD*), with *η* being the viscosity of water at 5°C.

### Production of full length hGHR

See Kassem et al.^48^ for expression, purification and reconstitution of hGHR in POPC-containing MSP1D1 nanodiscs.

### Phosphorus analysis

The POPC:GHR ratio of the formed nanodiscs with hGHR inserted was determined by phosphorus analysis^63^. This was done by hydrolyzing POPC in H_2_SO_4_ to release free phosphate (PO_4_^−3^), which reacted with molybdate to produce a blue chromophore, absorbing at 812 nm. A series of phosphate standards from 0 to 80 nM Na_2_HPO_4_ and hGHR in MSP1D1 at approximately 1 μM were prepared. Aliquots of 175 μL of each sample were transferred to glass tubes. HClO_4_ was added (400 μL 72 % (v/v)) to each sample and the glass tubes were loosely closed using glass pearls. The samples were heated to 180 °C in a water bath in a fume hood for 1 h and then left at room temperature to cool for 30 min. 4 mL of 125 mM (NH_4_)_6_Mo_7_O_24_ x 4 H_2_O was added to each sample and vortexed, followed by addition of 500 μL 10 % (w/w) ascorbic acid and vortexed again. Samples were then heated to 80 °C for 10 min in a water-bath and subsequently cooled in ice-water. Absorption was measured at 812 nm. A phosphate standard curve was generated, using the Na_2_HPO_4_ standards, by linear regression. The linear equation was then used to determine nmol content of phosphate in the hGHR in MSP1D1 samples. The ratio between POPC and hGHR(MSP1D1) was calculated from the ratios between their concentrations.

### Gel quantification of hGHR-loaded nanodiscs

Standards of hGHR and MSP1D1 with a known absorption at 280 nM were prepared and loaded in different amounts of the same gel as well as three aliquots of hGHR-loaded nanodiscs taken from three different positions of the SEC-elution profile (fraction 1, 2 and 3). The gels were stained with Coomassie brilliant blue G-250 (Bio-Rad) and subsequently destained in 15 % (v/v) ethanol, 5 % (v/v) acetic acid and 5 % glycerol (v/v). Gel images were obtained on a LAS4000 imager (GE Healthcare, USA) and the images were quantified in ImageJ^87^. The intensities of the standards were fitted by linear regression and the amount of hGHR relative to MSP1D1 quantified accordingly.

### Microscale thermophoresis

hGH was labelled with NT-647-NHS^88^ using the Monolith NT™ Protein Labelling Kit RED-NHS (NanoTemper Technologies) for 1 h at room temperature with NT-647-NHS at a molar ratio of 1:3 in labelling buffer following the protocol. These conditions favour the modification of the N-terminal amino group. Free dye was separated from reacted dye using the provided desalting column. The ratio between fluorophore and protein was 0.2. The equilibrium binding between 20 nM NT-647-NHS labelled hGH to hGHR(MSP1D1) was calculated from the change in thermophoresis Δ*F*_*norm*_ = Δ*F*_*hot*_/Δ*F*_*cold*_ measured on a Monolith NT.115 (NanoTemper Technologies). For hGH_G120R_ the raw fluorescence change was used to determine the binding affinity. A two-fold dilution series of monomeric hGHR from 750 nM to 23 pM was prepared in 20 mM Na_2_HPO4/NaH_2_PO_4_ (pH 7.4), 100 mM NaCl and measured in triplicates. Samples were loaded into Monolith NT.115 Premium Capillaries (NanoTemper Technologies), and thermophoresis and raw fluorescence signals measured at 25 °C with a light-emitting diode (LED) power of 80% and an infrared (IR) laser power of 100%. The dissociated constant *K_D_* was obtained by fitting the data by:

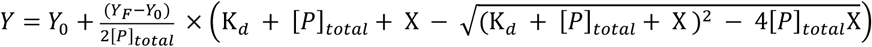

where *Y* is the measured fluorescence/MST, *X* is the ligand, *[P]*_*tota*l_ is the total concentration of the protein, *Y_F_* is the estimated end point of the titration and *Y_0_* is the start point.

### N-glycosylation removal by endoglycosidase H

1 μg purified full length hGHR was incubated with 500 units of Endo-H (New Biolabs, USA) at 4 °C in 20 mM NaH_2_PO_4_/Na_2_HPO_4_ (pH 7.4), 150 mM NaCl and 5 % (v/v) glycerol. The sample was separated and analysed on a 15 % SDS-PAGE gel and visualized by in-gel fluorescence on a LAS 4000 Imager (GE Healthcare, USA).

### Western blotting

hGHR was separated on a 15 % SDS-PAGE gel and blotted to a PDVF membrane as described in Pedersen et al., 1996^89^. Horse radish peroxidase conjugated Concanavalin-A (SigmaAldrich L6397) was used to identify O-glycosylations after western blotting. Chemiluminescence was detected by using the immobilon western chemiluminescent HRP substrate from Millipore ® and the LAS4000 imager (GE Healthcare, USA).

### Small angle x-ray and neutron scattering

SAXS data on hGH, hGHR-ECD and the hGH:hGHR-hECD 1:1 and 1:2 complexes were collected at the PETRA III, P12 beamline (DESY synchrotron, Hamburg)^90^, following standard procedures and at 8°C. All samples were concentrated and run on a Superdex 200 increase 10/300 GL in 20 mM Na_2_HPO_4_/NaH_2_PO_4_ pH 7.4, 150 mM NaCl prior to measuring. The most concentrated top fractions were taken, except for 1:1 complex, where the fraction was taken to the right of the peak, to make sure the hGH:hGHR-ECD 1:2 complex was absent in the sample. hGH was measured at 1.8 mg/mL, ECD at 3.5 mg/mL, hGH:hGHR-ECD 1:1 complex at 0.3 mg/mL and the hGH:hGHR-ECD 1:2 complex at 1.3 mg/mL. The scattering curves each of which is an average of 40 frames were recorded and the buffer was measured before and after each sample. Processing and preliminary analysis of data was done using the ATSAS package^91^. As a part of the process, buffer scattering curves before and after the sample were averaged and subtracted from the scattering curve of the sample. The scattering curves were scaled into units of 1/cm via the ATSAS package^91^ and using a measurement of water as secondary standard. The data was logarithmically re-binned. For the full length hGHR in MSP1D1, in-line SEC-SAXS of the sample in 20 mM Na_2_HPO_4_/NaH_2_PO_4_ pH 7.4, 150 mM NaCl was performed at BM29 (ESRF, Grenoble) equipped with a Superose 6 increase 10/300 GL (GE health care) running at a flow rate of 0.75 mL/min. In-line SEC-SANS data on the full length hGHR in MSP1D1 was recorded on the D22 small-angle scattering diffractometer at ILL, Grenoble, France. The in-line SEC was performed using a the recently commissioned and described modular HPLC system (Serlabo) in 20 mM Na_2_HPO_4_/NaH_2_PO_4_ pH 7.4, 150 mM NaCl on a Superose 6 increase 10/300 GL (GE health care)^61,92^. The flow rate was lowered from the 0.75 mL/min used in the SEC-SAXS measurements to 0.05 ml/min when the peak was reached in the lower intensity SANS to get as good counting statistics on the individual frames as possible. Two settings were used, 11.2 and 2.0 m (with collimation lengths of 11.2 and 2.8 m, respectively), giving a q-range between 0.0044–0.46 A°^−1^. The intensities were binned into 30 s frames.

### Modelling of the hGHR-ECD

To build a model of the full-length hGHR-ECD that covers the same sequence of the construct used in the experimental procedures the following steps were performed: i) An available structure of the GRH-ECD from the PDB was selected (chain C of 3HHR, residues 32 to 236). ii) Missing loops on the structure (57-61 and 74-77) were completed using the MODELLER^93^ interface of CHIMERA^94^. iii) The missing N-terminal (residues 1 to 31) and C-terminal (residues 237 to 245) tails were modelled as ensembles in order to capture their flexibility in the fitting of SAXS data. The ROSETTA ^95^ routine “Floppy tail” ^96,97^ was employed to generate 5000 conformations of both tails.

### Modelling of the hGHR ECD-TMD linker

The linker between the hGHR-ECD and hGHR-TMD (S237-W249) is not present in the available structures of hGHR-ECD and its structure may play a relevant role in determining the proper ECD-TMD orientation. Thus, this linker was modelled to provide a starting conformation of the hECD-TMD part of hGHR for further use in the modelling of the full-length hGHR structure.. To do this, the recently developed mp_domain_assembly protocol^98^ was implemented in Rosetta_MP^99^ was used. The structure of the ECD used correspond to chain C of 3HHR with completed loops (residues 32-236) and the TMD structure correspond to an NMR derived models (residues 250-272). A total of 5000 models were built with the best 10 (according to their Rosetta score) selected for further analysis, and the best ranked model used as a rigid body in the semi-analytical models of hGHR in a nanodisc and as starting conformation in the building of the full-length hGHR CG model (see below).

### Structural model for the full length hGHR-GFP

A full-length model of intact hGHR with a GFP on its C-terminus was built using the different pieces modelled separately. A representative conformation from the re-weighted sub-ensemble of the full-length ECD (residues 1-237) was aligned to the best model of the ECD-TMD to obtain a complete ECD-TMD structure (residues 1-272). A representative structure of the ICD (residues 273-620) was taken from the back-mapped conformation from the CG-MetaD-*R_g_* simulation with 10% increase in the protein-water interaction strength as described in *Results*. Rotations of the peptide bond between residues 273-274 had to be adjusted to allow the correct orientation of the ICD with respect to the TMD and the membrane plane. EGFP (PDB 1EMA) was added at residue 620. The all-atom model was used to build a CG system using the Martini Maker module^100^ of CHARMM-GUI^101^ to obtain a system of protein + POPC + water + 150 mM NaCl for the martini 2 forcefield. The topology was later adapted to open beta version of martini 3 (m3.b3.2)^102^. The final system contains 453662 beads and has a size of 361×361×406 Å^3^.

### Coarse grained MD simulations

All MD simulations were performed using Gromacs 2016 and 2018^103^ using the open beta version of the Martini 3 (3.b3.2) force field^102,104,105^ that was modified in order to avoid excessive compaction of the disordered regions. To find the optimal factor by which an increase in the protein water-interactions better reproduces the R_g_ of the hGHR-ICD, two sets of simulations were performed in which different values of the protein-water interaction strengths ranging between 5% to 15% were used. Unbiased simulations were performed with a 5%, 6%, 8% and 10% increase, while metadynamics simulations (see below) were performed with a 10%, 11%, 12%, 13%, 14% and 15% increase. Based on the best reproduction of *R_g_* (see **Fig. S3**) we chose a 10% increase in interaction strength and used this also for the simulation of the hGHR-GFP+POPC system. Simulation parameters were chosen following the recommendations in^106^. Briefly, the Verlet cut-off scheme was considering a buffer tolerance of 0.005 kJ/(mol ps atom) The reaction-field method was used for Coulomb interactions with a cut-off of 11 Å and a dielectric constant of *εr* = 15 for water. For van der Waals interactions the cut-off scheme with a cut-off of 11 Å was used. The velocity rescaling thermostat was employed with a reference temperature of T = 300 K and 310 K for the hGHR-ICD and hGHR-GFP+POPC simulations respectively, with a coupling constant of *τ*T = 1 ps^107^ in all cases. For the equilibrations, the Berendsen barostat was employed *(p* = 1 bar, *τp* = 3 ps), whereas the production runs were performed with a Parrinello-Rahman barostat (p = 1 bar, *τp* = 12 ps)^108^. A semi-isotropic pressure coupling was used for the hGHR system embedded on a lipid bilayer. For all systems an initial round of equilibrations with decreasing constraints applied to the protein beads (hGHR-ICD) and protein beads and lipid beads (hGHR-GFP) was performed.

Sampling of the hGHR-ICD simulations with an increase in the protein-water interactions of 10%, 11%, 12%, 13%, 14% and 15% was enhanced using a well-tempered metadynamics^109^ protocol applied with PLUMED 2.5^110^. The *R_g_* of the protein was used as collective variable (CV) within the boundaries of 30 to 110 Å. The metadynamics parameters used are: a bias factor of 50, gaussian height of 4.2 kJ/mol and collective variable space Gaussian widths equal to 0.3.

Analysis of the MD trajectories was performed using plugins and analysis tools implemented in VMD^111^, GROMACS and PLUMED together with in-house prepared tcl and python scripts. All molecular renderings were done with VMD.

### Fitting of the SAXS data of the hGHR-ECD and hGHR-ICD by the MD models

Similar protocols were utilized to fit the SAXS data of the hGHR-ECD and hGHR-ICD with the conformations obtained from the modelling of hGHR-ECD and MD simulations of hGHR-ICD, respectively: i) For the hGHR-ECD the SAXS profile of each conformation was directly calculated and fitted to the SAXS data using Pepsi-SAXS^112^ with all parameters free. For the conformations obtained from the different hGHR-ICD simulations, an initial round of back-mapping was performed to go from a coarse-grained to all-atoms as described in^58^, before calculating and fitting its SAXS profile with Pepsi-SAXS. ii) From the fits, the average value of the hydration shell contrast was calculated (hGHR-ECD =7.4%; hGHR-ICD = 4%) and used as a fixed parameter in a second round of fitting. iii) The average scattering profile of the ensemble was calculated and compared to the data. iv) In the case of hGHR-ECD, the BME^41,42^ procedure, designed to integrate ensembles of molecular models (and simulations) with experiments was used to reweight the ensemble against the experimental data and refine the fitting. From the reweighted ensemble a representative sub-ensemble of 500 conformations was obtained

### Semi-analytical model for the ND embedded hGHR-GFP

To generate the semi-analytical model for the full-length hGHR, a combination of analytical approaches to describe the nanodisc and the hGHR-ICD, and rigid body modelling for the ECD-TMD and GFP was implemented the *WillItFit*^70^ framework. The mathematical model for hGHR in nanodiscs, illustrated in Fig. 5A, is composed by four distinct amplitude components arising from the ECD-TMD, the ICD, the attached GFP and the surrounding nanodisc. The final expression for the total scattering intensity was calculated on absolute scale as the scattering amplitude squared:

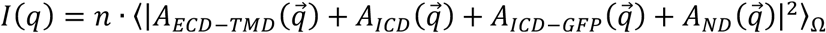

where 〈… 〉_Ω_ denote the orientational average, |… | denote the complex norm, *n* is the number-density of particles and 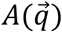 is the scattering amplitude of each component for a single particle. Subscript *ECD-TMD* refers to the hGHR-ECD with transmembrane domain, *ICD* refers to the intrinsically disordered intracellular domain and *ICD-GFP* refers to the GFP fused to the ICD and *ND* refers to the POPC loaded nanodisc. For each amplitude term, 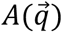, we furthermore have that 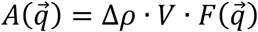, where *Δρ* is the average excess scattering length density, *V* is the volume and 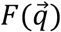 is the normalized form factor amplitude for the relevant component. The model for 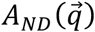 is the same as we have described previously^69^: A stack of five elliptical cylinders representing the phospholipid bilayer was surrounded by a hollow elliptical cylinder representing the two stacked MSP’s. As usual^69^ molecular constraints were systematically implemented to constrain the solution space. As a part of this, the height of the MSP was fixed to a value of 25.8 Å as derived from a high-resolution structure of nanodiscs^113^. The scattering amplitudes of the hGHR-ECD-TMD and the GFP were calculated from their atomic coordinates as a part of our *WillItFit*^70^ framework and as outlined in previous work^37^ and incorporated into the ND rigid bodies. PDB 1EMA was used for the GFP atomic coordinates while those of the flexible hGHR-ECD-TMD were represented by a single of the structures obtained from the modelling of the ECD-TMD linker with the N-terminal tail added from the ensemble produced in the modeling of the full-length hGHR-ECD as described in previous sections. This allowed the TMD to displace lipids in the ND and for adjusting the excess scattering lengths of the lipid embedded residues by considering their lipid environment rather than the solvent^37^. The averaged form factor intensity for the hGHR-ICD was modeled as a Gaussian Random Coil through the Debye function^51^. The averaged form factor amplitude for a Gaussian random coil required for the cross-terms in the calculation of *I(q)* is given by the so-called Hammouda function^114^ which is a function of the ensemble average *R_g_* of the coil. Hence, we used the same modelling principle as previously applied for polymer modified micelles^115^ to connect the hGHR-ICD to the nanodisc embedded TMD in the model. Following a similar philosophy, the GFP was randomly oriented and located within a certain allocated “confusion volume”. This way the model captures the dynamically evolving position of the GFP with respect to the rest of the system. For the modelling of the confusion volume we attempted to mimic the bowl-like distribution of GFP below the bilayer as observed in the CG-MD simulation of hGHR in a lipid bilayer (see Fig 6F), by placing the GFP randomly in a thick cylindrical shell below the nanodisc (see Fig 6A, inner and outer shell radii equal to, respectively, 1 and 1.5 times the *R*_g_ of hGHR-ICD). However, we found that the actual shape of the confusion volume, whether it was bowl-shaped or simply spherical, only had a minor effect.

## Footnotes

1 Of note, in these simulations, we did not consider the formation of transient secondary structures previously observed by NMR^16^. However, at the resolution provided by SAXS this is a reasonable approach, which was also applied in the modeling and simulation of the full-length hGHR.

2 Due to the low signal at the right side of the SEC-SANS peak, the data from the monomeric fractions were too noisy to allow for a robust further analysis.

## Notes

### Competing Interest Statement

The authors have declared no competing interest.

## REFERENCES

1. Madsen, K., Friberg, U., Roos, P., Edén, S. & Isaksson, O. Growth hormone stimulates the proliferation of cultured chondrocytes from rabbit ear and rat rib growth cartilage. Nature 304, 545–7 (2005).

2. Lupu, F., Terwilliger, J. D., Lee, K., Segre, G. V. & Efstratiadis, A. Roles of growth hormone and insulin-like growth factor 1 in mouse postnatal growth. Dev. Biol. 229, 141–62 (2001).

3. Waters, M. J. & Brooks, A. J. Growth hormone receptor: Structure function relationships. Horm. Res. Paediatr. 76, 12–16 (2011).

4. Isaksson, O. G., Jansson, J. O. & Gause, I. A. Growth hormone stimulates longitudinal bone growth directly. Science 216, 1237–9 (1982).

5. Guler, H. P., Zapf, J., Scheiwiller, E. & Froesch, E. R. Recombinant human insulin-like growth factor I stimulates growth and has distinct effects on organ size in hypophysectomized rats. Proc. Natl. Acad. Sci. U. S. A. 85, 4889–93 (1988).

6. Yakar, S. et al. Circulating levels of IGF-1 directly regulate bone growth and density. J. Clin. Invest. 110, 771–81 (2002).

7. Chhabra, Y. et al. A growth hormone receptor SNP promotes lung cancer by impairment of SOCS2-mediated degradation. Oncogene 37, 489–501 (2018).

8. Chanson, P. & Salenave, S. Acromegaly. Orphanet J. Rare Dis. 3, 17 (2008).

9. Melmed, S. et al. Current status and future opportunities for controlling acromegaly. Pituitary 5, 185–96 (2002).

10. Duncan, E. & Wass, J. A. Investigation protocol: acromegaly and its investigation. Clin. Endocrinol. 50, 285–93 (1999).

11. Mullis, P. E. Genetics of isolated growth hormone deficiency. J. Clin. Res. Pediatr. Endocrinol. 2, 52–62 (2010).

12. Reh, C. S. & Geffner, M. E. Somatotropin in the treatment of growth hormone deficiency and Turner syndrome in pediatric patients: A review. Clin. Pharmacol. Adv. Appl. 2, 111–122 (2010).

13. Shimatsu, A., Nagashima, M., Hashigaki, S., Ohki, N. & Chihara, K. Efficacy and safety of monotherapy by pegvisomant, a growth hormone receptor antagonist, in Japanese patients with acromegaly. Endocr. J. 63, 337–47 (2016).

14. Brooks, A. J., Dehkhoda, F. & Kragelund, B. B. Cytokine Receptors. in Principles of Endocrinology and Hormone Action (eds. Belfiore, A. & LeRoith, D.) 157–185 (Springer International Publishing, 2018). doi:10.1007/978-3-319-44675-2_8

15. Seiffert, P. et al. Orchestration of signaling by structural disorder in class 1 cytokine receptors. Cell Commun. Signal. resubmitted, (2020).

16. Haxholm, G. W. et al. Intrinsically disordered cytoplasmic domains of two cytokine receptors mediate conserved interactions with membranes. Biochem. J. 468, 495–506 (2015).

17. Waters, M. J. The growth hormone receptor. Growth Horm. IGF Res. 28, 6–10 (2016).

18. Liongue, C. & Ward, A. C. Evolution of Class I cytokine receptors. BMC Evol. Biol. 7, 120 (2007).

19. Baumgartner, J. W., Wells, C. A., Chen, C. M. & Waters, M. J. The role of the WSXWS equivalent motif in growth hormone receptor function. J. Biol. Chem. 269, 29094–101 (1994).

20. van den Eijnden, M. J. M., Lahaye, L. L. & Strous, G. J. Disulfide bonds determine growth hormone receptor folding, dimerisation and ligand binding. J. Cell Sci. 119, 3078–86 (2006).

21. Olsen, J. G. & Kragelund, B. B. Who climbs the tryptophan ladder? On the structure and function of the WSXWS motif in cytokine receptors and thrombospondin repeats. Cytokine Growth Factor Rev. 25, 337–341 (2014).

22. de Vos, A. M., Ultsch, M. & Kossiakoff, A. A. Human growth hormone and extracellular domain of its receptor: crystal structure of the complex. Science 255, 306–12 (1992).

23. Brooks, A. J. et al. Mechanism of activation of protein kinase JAK2 by the growth hormone receptor. Science 344, 1249783 (2014).

24. Wilmes, S. et al. Mechanism of homodimeric cytokine receptor activation and dysregulation by oncogenic mutations. Science 367, 643–652 (2020).

25. Brown, R. J. et al. Model for growth hormone receptor activation based on subunit rotation within a receptor dimer. Nat. Struct. Mol. Biol. 12, 814–821 (2005).

26. Fuh, G. et al. Rational design of potent antagonists to the human growth hormone receptor. Science 256, 1677–80 (1992).

27. Sundström, M. et al. Crystal structure of an antagonist mutant of human growth hormone, G120R, in complex with its receptor at 2.9 Å resolution. J. Biol. Chem. 271, 32197–32203 (1996).

28. Chantalat, L., Jones, N. D., Korber, F., Navaza, J. & Pavlovsky, A. G. THE CRYSTAL-STRUCTURE OF WILD-TYPE GROWTH-HORMONE AT 2.5 ANGSTROM RESOLUTION. Protein Pept.Lett. 2, 333–340 (1995).

29. Bocharov, E. V. et al. Structural basis of the signal transduction via transmembrane domain of the human growth hormone receptor. Biochim. Biophys. Acta - Gen. Subj. 1862, 1410–1420 (2018).

30. Bugge, K. et al. A combined computational and structural model of the full-length human prolactin receptor. Nat. Commun. 7, 1–11 (2016).

31. Carroni, M. & Saibil, H. R. Cryo electron microscopy to determine the structure of macromolecular complexes. Methods 95, 78–85 (2016).

32. Kassem, N., Kassem, M. M., Pedersen, S. F., Pedersen, P. A. & Kragelund, B. B. Yeast recombinant production of intact human membrane proteins with long intrinsically disordered intracellular regions for structural studies. Biochim. Biophys. acta. Biomembr. 1862, 183272 (2020).

33. Breyton, C. et al. Small angle neutron scattering for the study of solubilised membrane proteins. Eur. Phys. J. E. Soft Matter 36, 71 (2013).

34. Svergun, D. I. & Koch, M. H. J. Reports on Progress in Physics Related content Small-angle scattering studies of biological macromolecules in solution Small-angle scattering studies of biological macromolecules in solution. Reports Prog. Phys. 1735 (2003).

35. Bayburt, T. H., Grinkova, Y. V. & Sligar, S. G. Self-Assembly of Discoidal Phospholipid Bilayer Nanoparticles with Membrane Scaffold Proteins. Nano Lett. 2, 853–856 (2002).

36. Skar-Gislinge, N. et al. Small-angle scattering determination of the shape and localization of human cytochrome P450 embedded in a phospholipid nanodisc environment. Acta Crystallogr. D. Biol. Crystallogr. 71, 2412–21 (2015).

37. Kynde, S. A. R. et al. Small-angle scattering gives direct structural information about a membrane protein inside a lipid environment. Acta Crystallogr. Sect. D Biol. Crystallogr. 70, 371–383 (2014).

38. Bugge, K., Lindorff-Larsen, K. & Kragelund, B. B. Understanding single-pass transmembrane receptor signaling from a structural viewpoint-what are we missing? FEBS J. 283, 4424–4451 (2016).

39. Dagil, R. et al. The WSXWS motif in cytokine receptors is a molecular switch involved in receptor activation: Insight from structures of the prolactin receptor. Structure 20, 270–282 (2012).

40. Glatter, O. A new method for the evaluation of small-angle scattering data. J. Appl. Crystallogr. 10, 415–421 (1977).

41. Ahmed, M. C., Crehuet, R. & Lindorff-Larsen, K. Analyzing and comparing the radius of gyration and hydrodynamic radius in conformational ensembles of intrinsically disordered proteins. bioRxiv 679373 (2019). doi:10.1101/679373

42. Bottaro, S., Bengtsen, T. & Lindorff-Larsen, K. Integrating Molecular Simulation and Experimental Data: A Bayesian/Maximum Entropy Reweighting Approach. bioRxiv 457952 (2018). doi:10.1101/457952

43. Bugge, K., Steinocher, H., Brooks, A. J., Lindorff-Larsen, K. & Kragelund, B. B. Exploiting hydrophobicity for efficient production of transmembrane helices for structure determination by NMR spectroscopy. Anal. Chem. 87, 9126–31 (2015).

44. Shen, Y. & Bax, A. Identification of helix capping and b-turn motifs from NMR chemical shifts. J. Biomol. NMR 52, 211–32 (2012).

45. Güntert, P. Automated NMR structure calculation with CYANA. Methods Mol. Biol. 278, 353–78 (2004).

46. Khondker, A. et al. Membrane charge and lipid packing determine polymyxin-induced membrane damage. Commun. Biol. 2, 1–11 (2019).

47. Bürck, J., Wadhwani, P., Fanghänel, S. & Ulrich, A. S. Oriented Circular Dichroism: A Method to Characterize Membrane-Active Peptides in Oriented Lipid Bilayers. Acc. Chem. Res. 49, 184–92 (2016).

48. Kassem, N., Kassem, M. M., Pedersen, S. F., Pedersen, P. A. & Kragelund, B. B. Yeast recombinant production of intact human membrane proteins with long intrinsically disordered intracellular regions for structural studies. BBA - Biomembr. 1862(6), 183272 (2019).

49. Brown, W. & Mortensen, K. Comparison of Correlation Lengths in Semidilute Polystyrene Solutions in Good Solvents by Quasi-Elastic Light Scattering and Small-Angle Neutron Scattering. Macromolecules 21, 420–425 (1988).

50. Flory, P. J. Principles of Polymer Chemistry. Cornless University press (1953).

51. Debye, P. Molecular-weight determination by light scattering. J. Phys. Colloid Chem. 51, 18–32 (1947).

52. Kohn, J. E. et al. Random-coil behavior and the dimensions of chemically unfolded proteins. Proc. Natl. Acad. Sci. U. S. A. 101, 12491–6 (2004).

53. Zheng, W. & Best, R. B. An Extended Guinier Analysis for Intrinsically Disordered Proteins. J. Mol. Biol. 430, 2540–2553 (2018).

54. Riback, J. A. et al. Innovative scattering analysis shows that hydrophobic disordered proteins are expanded in water. Science 358, 238–241 (2017).

55. Marsh, J. A. & Forman-kay, J. D. Sequence Determinants of Compaction in Intrinsically Disordered Proteins. Biophys J 98, 2383–2390 (2010).

56. Stark, A. C., Andrews, C. T. & Elcock, A. H. Toward optimized potential functions for protein-protein interactions in aqueous solutions: osmotic second virial coefficient calculations using the MARTINI coarse-grained force field. J. Chem. Theory Comput. 9, 4176–4185 (2013).

57. Javanainen, M., Martinez-Seara, H. & Vattulainen, I. Excessive aggregation of membrane proteins in the Martini model. PLoS One 12, e0187936 (2017).

58. Larsen, A. H. et al. Combining molecular dynamics simulations with small-angle X-ray and neutron scattering data to study multi-domain proteins in solution. PLoS Comput. Biol. 16, e1007870 (2020).

59. Milkovic, N. M. et al. Interplay of folded domains and the disordered low-complexity domain in mediating hnRNPA1 phase separation. bioRxiv (2020). doi:10.1101/2020.05.15.096966

60. Skar-Gislinge, N., Johansen, N. T., Høiberg-Nielsen, R. & Arleth, L. Comprehensive Study of the Self-Assembly of Phospholipid Nanodiscs: What Determines Their Shape and Stoichiometry? Langmuir 34, 12569–12582 (2018).

61. Johansen, N. T., Pedersen, M. C., Porcar, L., Martel, A. & Arleth, L. Introducing SEC-SANS for studies of complex self-organized biological systems. Acta Crystallogr. Sect. D, Struct. Biol. 74, 1178–1191 (2018).

62. Gent, J., van Kerkhof, P., Roza, M., Bu, G. & Strous, G. J. Ligand-independent growth hormone receptor dimerization occurs in the endoplasmic reticulum and is required for ubiquitin system-dependent endocytosis. Proc. Natl. Acad. Sci. U. S. A. 99, 9858–63 (2002).

63. Rouser, G., Siakotos, A. N. & Fleischer, S. Quantitative analysis of phospholipids by thin-layer chromatography and phosphorus analysis of spots. Lipids 1, 85–6 (1966).

64. Harding, P. A. et al. In vitro mutagenesis of growth hormone receptor Asn-linked glycosylation sites. Mol. Cell. Endocrinol. 106, 171–80 (1994).

65. Bjørkskov, F. B. et al. Purification and functional comparison of nine human Aquaporins produced in Saccharomyces cerevisiae for the purpose of biophysical characterization. Sci. Rep. 7, 1–21 (2017).

66. Wennbo, H. et al. Activation of the prolactin receptor but not the growth hormone receptor is important for induction of mammary tumors in transgenic mice. J. Clin. Invest. 100, 2744–51 (1997).

67. Bernat, B., Sun, M., Dwyer, M., Feldkamp, M. & Kossiakoff, A. A. Dissecting the binding energy epitope of a high-affinity variant of human growth hormone: cooperative and additive effects from combining mutations from independently selected phage display mutagenesis libraries. Biochemistry 43, 6076–84 (2004).

68. Kouadio, J.-L. K., Horn, J. R., Pal, G. & Kossiakoff, A. A. Shotgun alanine scanning shows that growth hormone can bind productively to its receptor through a drastically minimized interface. J. Biol. Chem. 280, 25524–32 (2005).

69. Skar-Gislinge, N. et al. Elliptical structure of phospholipid bilayer nanodiscs encapsulated by scaffold proteins: casting the roles of the lipids and the protein. J. Am. Chem. Soc. 132, 13713–22 (2010).

70. Pedersen, M. C., Arleth, L. & Mortensen, K. WillItFit: a framework for fitting of constrained models to small-angle scattering data. J. Appl. Crystallogr. 46, 1894–1898 (2013).

71. Minezaki, Y., Homma, K. & Nishikawa, K. Intrinsically disordered regions of human plasma membrane proteins preferentially occur in the cytoplasmic segment. J. Mol. Biol. 368, 902–13 (2007).

72. Kjaergaard, M. & Kragelund, B. B. Functions of intrinsic disorder in transmembrane proteins. Cell. Mol. Life Sci. 74, 3205–3224 (2017).

73. Ben-Avraham, D. et al. The GH receptor exon 3 deletion is a marker of male-specific exceptional longevity associated with increased GH sensitivity and taller stature. Sci. Adv. 3, e1602025 (2017).

74. Gouw, M. et al. The eukaryotic linear motif resource - 2018 update. Nucleic Acids Res. 46, D428–D434 (2018).

75. Hamming, O. J. et al. Crystal structure of interleukin-21 receptor (IL-21R) bound to IL-21 reveals that sugar chain interacting with WSXWS motif is integral part of IL-21R. J. Biol. Chem. 287, 9454–9460 (2012).

76. Hofsteenge, J. et al. New Type of Linkage between a Carbohydrate and a Protein: C-Glycosylation of a Specific Tryptophan Residue in Human RNase Us. Biochemistry 33, 13524–13530 (1994).

77. Bürgi, J., Xue, B., Uversky, V. N. & van der Goot, F. G. Intrinsic Disorder in Transmembrane Proteins: Roles in Signaling and Topology Prediction. PLoS One 11, e0158594 (2016).

78. Kaplan, M. et al. EGFR Dynamics Change during Activation in Native Membranes as Revealed by NMR. Cell 167, 1241–1251.e11 (2016).

79. Wang, X., Darus, C. J., Xu, B. C. & Kopchick, J. J. Identification of growth hormone receptor (GHR) tyrosine residues required for GHR phosphorylation and JAK2 and STAT5 activation. Mol. Endocrinol. 10, 1249–60 (1996).

80. Cormack, B. P. et al. Yeast-enhanced green fluorescent protein (yEGFP): A reporter of gene expression in Candida albicans. Microbiology 143, 303–311 (1997).

81. Khondker, A., Malenfant, D. J., Dhaliwal, A. K. & Rheinstädter, M. C. Carbapenems and Lipid Bilayers: Localization, Partitioning, and Energetics. ACS Infect. Dis. 4, 926–935 (2018).

82. Himbert, S. et al. Hybrid Erythrocyte Liposomes: Functionalized Red Blood Cell Membranes for Molecule Encapsulation. Adv. Biosyst. 4, 1–11 (2020).

83. Delaglio, F. et al. NMRPipe: A multidimensional spectral processing system based on UNIX pipes. J. Biomol. NMR 6, 277–293 (1995).

84. Vranken, W. F. et al. The CCPN data model for NMR spectroscopy: development of a software pipeline. Proteins 59, 687–96 (2005).

85. Shen, Y., Delaglio, F., Cornilescu, G. & Bax, A. TALOS+: a hybrid method for predicting protein backbone torsion angles from NMR chemical shifts. J. Biomol. NMR 44, 213–23 (2009).

86. Carper, W. R. Direct determination of quadrupolar and dipolar NMR correlation times from spin - lattice and spin - spin relaxation rates. Concepts Magn. Reson. 11, 51–60 (1999).

87. Schneider, C. A., Rasband, W. S. & Eliceiri, K. W. NIH Image to ImageJ: 25 years of image analysis. Nat. Methods 9, 671–5 (2012).

88. Jerabek-Willemsen, M. et al. MicroScale Thermophoresis: Interaction analysis and beyond. J. Mol. Struct. 1077, 101–113 (2014).

89. Pedersen, P. A., Rasmussen, J. H. & Joorgensen, P. L. Expression in high yield of pig alpha 1 beta 1 Na,K-ATPase and inactive mutants D369N and D807N in Saccharomyces cerevisiae. J Biol Chem 271, 2514–2522 (1996).

90. Blanchet, C. E. et al. Versatile sample environments and automation for biological solution X-ray scattering experiments at the P12 beamline (PETRA III, DESY). J. Appl. Crystallogr. 48, 431–443 (2015).

91. Petoukhov, M. V. et al. New developments in the ATSAS program package for small-angle scattering data analysis. J. Appl. Crystallogr. 45, 342–350 (2012).

92. Jordan, A. et al. SEC-SANS: size exclusion chromatography combined in situ with small-angle neutron scattering. J. Appl. Crystallogr. 49, 2015–2020 (2016).

93. Webb, B. & Sali, A. Protein Structure Modeling with MODELLER. in 1–15 (2014). doi:10.1007/978-1-4939-0366-5_1

94. Pettersen, E. F. et al. UCSF Chimera--a visualization system for exploratory research and analysis. J. Comput. Chem. 25, 1605–12 (2004).

95. Leaver-Fay, A. et al. ROSETTA3: an object-oriented software suite for the simulation and design of macromolecules. Methods Enzymol. 487, 545–74 (2011).

96. Kleiger, G., Saha, A., Lewis, S., Kuhlman, B. & Deshaies, R. J. Rapid E2-E3 assembly and disassembly enable processive ubiquitylation of cullin-RING ubiquitin ligase substrates. Cell 139, 957–68 (2009).

97. Zhang, J., Lewis, S. M., Kuhlman, B. & Lee, A. L. Supertertiary structure of the MAGUK core from PSD-95. Structure 21, 402–13 (2013).

98. Koehler Leman, J. & Bonneau, R. A Novel Domain Assembly Routine for Creating Full-Length Models of Membrane Proteins from Known Domain Structures. Biochemistry 57, 1939–1944 (2018).

99. Alford, R. F. et al. An Integrated Framework Advancing Membrane Protein Modeling and Design. PLoS Comput. Biol. 11, e1004398 (2015).

100. Qi, Y. et al. CHARMM-GUI Martini Maker for Coarse-Grained Simulations with the Martini Force Field. J. Chem. Theory Comput. 11, 4486–94 (2015).

101. Jo, S., Kim, T., Iyer, V. G. & Im, W. CHARMM-GUI: a web-based graphical user interface for CHARMM. J. Comput. Chem. 29, 1859–65 (2008).

102. CG martini. Available at: http://www.cgmartini.nl/index.php/force-field-parameters/particle-definitions.

103. Abraham, M. J. et al. GROMACS: High performance molecular simulations through multi-level parallelism from laptops to supercomputers. SoftwareX 1–2, 19–25 (2015).

104. Marrink, S. J. & Tieleman, D. P. Perspective on the Martini model. Chem. Soc. Rev. 42, 6801–22 (2013).

105. Alessandri, R. et al. Pitfalls of the Martini Model. J. Chem. Theory Comput. 15, 5448–5460 (2019).

106. de Jong, D. H., Baoukina, S., Ingólfsson, H. I. & Marrink, S. J. Martini straight: Boosting performance using a shorter cutoff and GPUs. Comput. Phys. Commun. 199, 1–7 (2016).

107. Bussi, G., Donadio, D. & Parrinello, M. Canonical sampling through velocity rescaling. J. Chem. Phys. 126, 014101 (2007).

108. Parrinello, M. & Rahman, A. Polymorphic transitions in single crystals: A new molecular dynamics method. J. Appl. Phys. 52, 7182–7190 (1981).

109. Barducci, A., Bussi, G. & Parrinello, M. Well-tempered metadynamics: a smoothly converging and tunable free-energy method. Phys. Rev. Lett. 100, 020603 (2008).

110. Granseth, E., Seppälä, S., Rapp, M., Daley, D. O. & Von Heijne, G. Membrane protein structural biology - How far can the bugs take us? (Review). Mol. Membr. Biol. 24, 329–332 (2007).

111. Humphrey, W., Dalke, A. & Schulten, K. VMD: visual molecular dynamics. J. Mol. Graph. 14, 33–8, 27–8 (1996).

112. Grudinin, S., Garkavenko, M. & Kazennov, A. Pepsi-SAXS: an adaptive method for rapid and accurate computation of small-angle X-ray scattering profiles. Acta Crystallogr. Sect. D, Struct. Biol. 73, 449–464 (2017).

113. Bibow, S. et al. Solution structure of discoidal high-density lipoprotein particles with a shortened apolipoprotein A-I. Nat. Struct. Mol. Biol. 24, 187–193 (2017).

114. Hammouda, B. Structure factor for starburst dendrimers. J. Polym. Sci. Part B Polym. Phys. 30, 1387–1390 (1992).

115. Pedersen, J. S. Form factors of block copolymer micelles with spherical, ellipsoidal and cylindrical cores. J. Appl. Crystallogr. 33, 637–640 (2000).

116. Sonnhammer, E. L., von Heijne, G. & Krogh, A. A hidden Markov model for predicting transmembrane helices in protein sequences. Proceedings. Int. Conf. Intell. Syst. Mol. Biol. 6, 175–82 (1998).

117. Käll, L., Krogh, A. & Sonnhammer, E. L. L. A combined transmembrane topology and signal peptide prediction method. J. Mol. Biol. 338, 1027–36 (2004).

118. Käll, L., Krogh, A. & Sonnhammer, E. L. L. Advantages of combined transmembrane topology and signal peptide prediction--the Phobius web server. Nucleic Acids Res. 35, W429-32 (2007).

119. Jones, D. T. Improving the accuracy of transmembrane protein topology prediction using evolutionary information. Bioinformatics 23, 538–44 (2007).

120. Eisenberg, D., Schwarz, E., Komaromy, M. & Wall, R. Analysis of membrane and surface protein sequences with the hydrophobic moment plot. J. Mol. Biol. 179, 125–42 (1984).

121. UniProt Consortium. UniProt: a hub for protein information. Nucleic Acids Res. 43, D204-12 (2015).

