## Supplementary material for "Order and disorder – an integrative structure of the full-length human growth hormone receptor"

**List of supplementary figures:**
Supplementary Fig. S1: Analytical size exclusion chromatography of hGHR-ECD complexes
Supplementary Fig. S2: NMR analysis of the hGHR-TMD
Supplementary Fig. S3: Calibration of the protein-water interaction strength using Martini3 and MetaDynamics
Supplementary Fig. S4: Biochemical and biophysical analyses of hGHR in POPC-loaded nanodiscs
Supplementary Fig. S5: Additional insights of hGHR-GFP relative orientations from the CG-MD simulation

**List of supplementary tables:**
Supplementary Table S1: Molecular weights of the hGHR-ECD and ICD from the SAXS data
Supplementary Table S2: Dimensions of hGHR-ICD – comparisons of different analysis approaches
Supplementary Table S3: SAXS-refined fit parameters of the hGHR-GFP in nanodisc

**List of supplementary text:**
Detailed account of the results of the semi-analytical model fit of hGHR-GFP in POPC nanodiscs

**List of supplementary movies:**
Supplementary movie M1: Movie showing the 21 µs CG-MD simulation of hGHR-GFP in a POPC bilayer. Frames taken every 10 ns. Color and representation scheme as in Fig. 6.

### Supplementary Figures

Supplementary Fig. S1

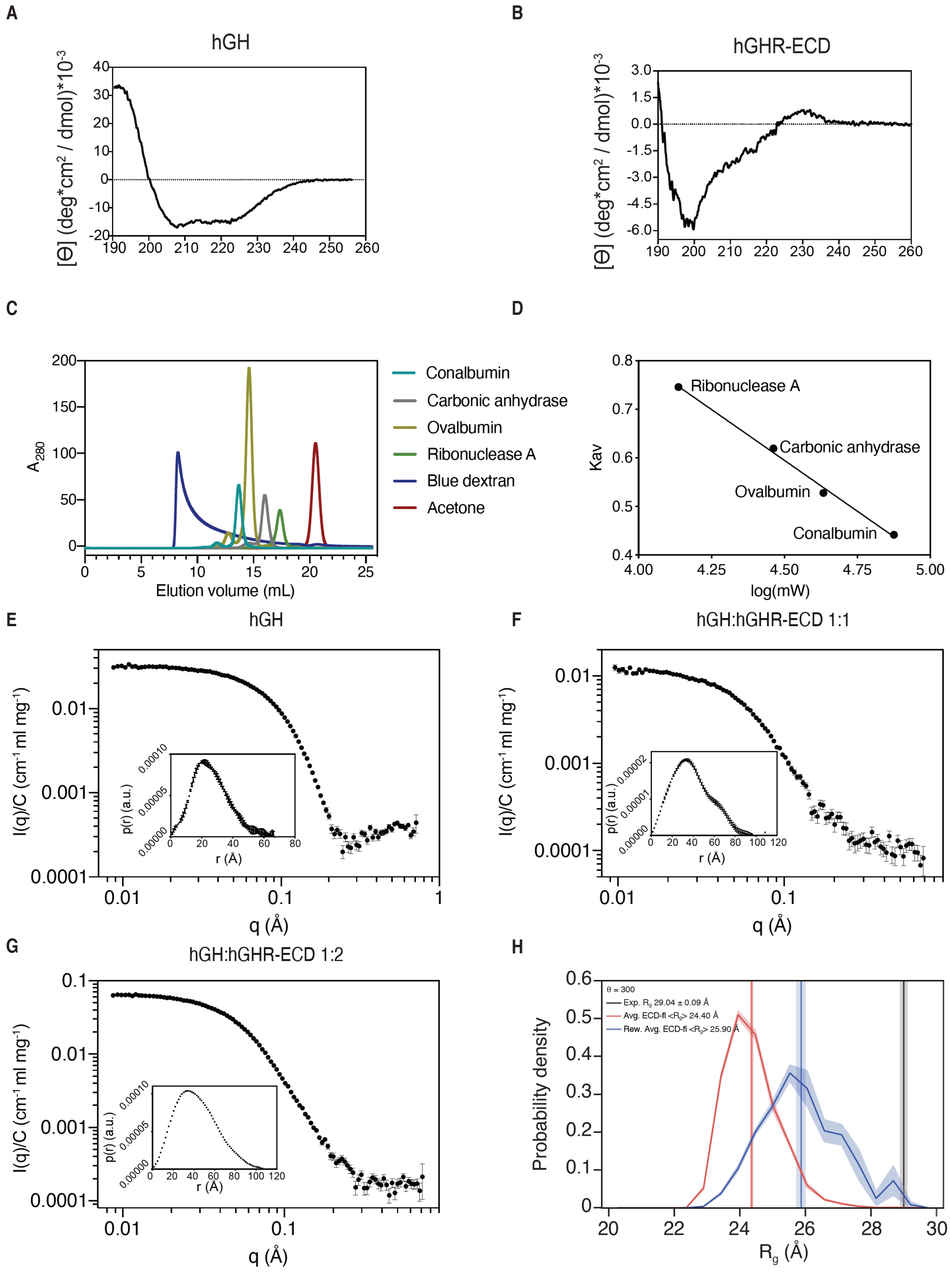


**Suppl. Fig. S1: Analytical size exclusion chromatography of hGHR-ECD complexes.** (A) Far-UV CD spectrum of hGH with a helicity calculated to be 32%[1] and 38%[2] (44% from the crystal structure PDB 1HGU). (B) Far-UV-CD spectrum of hGHR-ECD. (C) Six proteins standards; conalbumin, carbonic anhydrase, ovalbumin, ribonuclease A, blue dextran and acetone were run on a Superdex 200 increase 10/300 (GE Healthcare). (D) Partition coefficients were calculated for ribonuclease A, carbonic anhydrase, ovalbumin and conalbumin. Linear regression was performed, where $Kav = -0.4187*\log(MW)+2.4779$, $R^2 = 0.98$. SAXS curves of (E) hGH, (F) hGH:hGHR-ECD 1:1, and (G) hGH:hGHR-ECD 1:2. (H) $R_g$ distribution of hGHR-ECD models before (red) and after (blue) reweighting with BME. The vertical lines indicate the experimental $R_g$ values obtained from hGHR-ECD SAXS data (black) and the ensemble average $R_g$ from the hGHR-ECD models before (red) and after reweighting (blue). We note that the average (dry) $R_\mathrm{g}$ of 26 Å, is in good agreement with the experimental value of 29 Å, considering an expected 5-10% difference due to solvation shell scattering in the experiments.

Supplementary Fig. S2

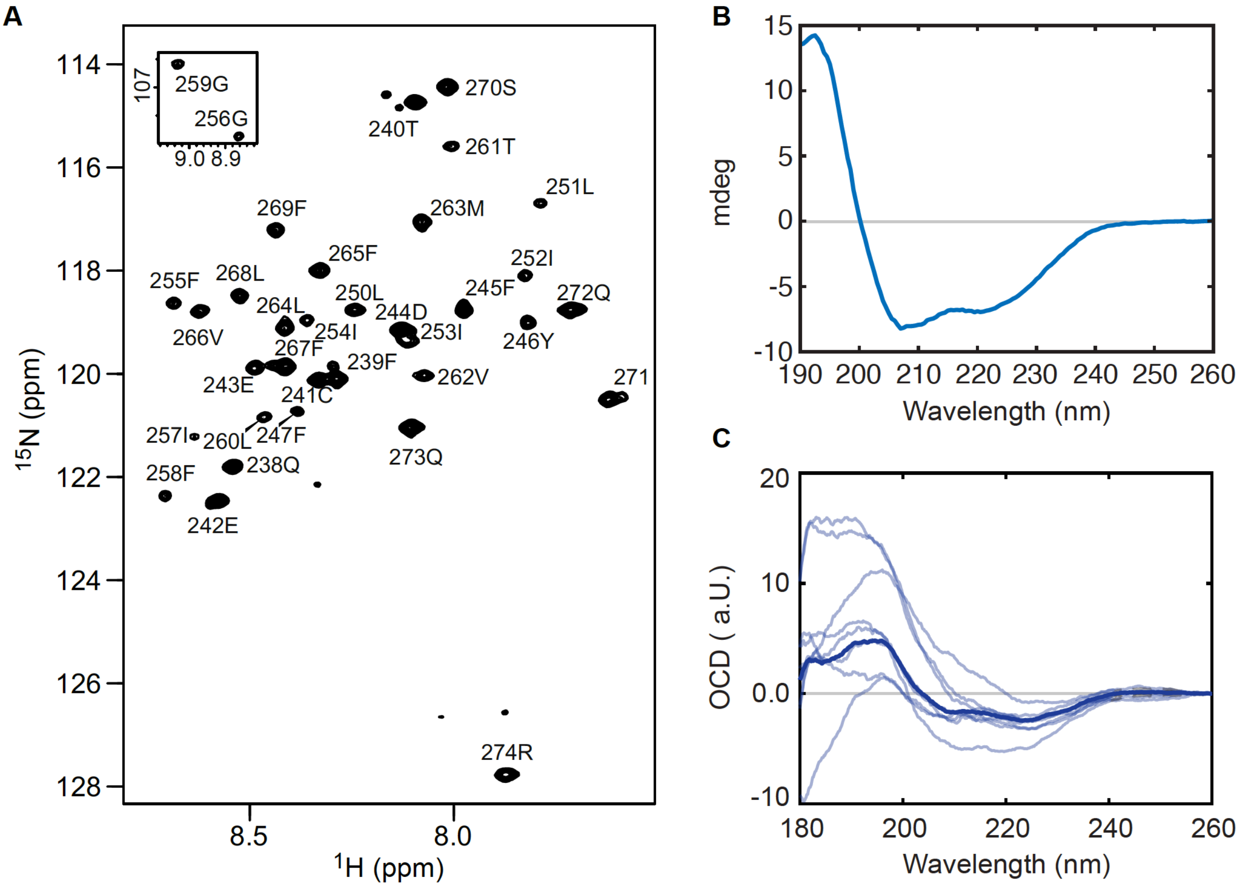


**Suppl. Fig. S2: Structural analysis of hGHR-TMD.** (A) Assigned $^{1}$H,$^{15}$N-HSQC spectrum at 37ºC of 1 mM $^{13}$C,$^{15}$N-hGHR-TMD in 210 mM DHPC. (B) Far-UV CD spectrum of 10 µM hGHR-TMD in 2 mM DHPC. (C) OCD spectra of 6 µg hGHR-TMD in 50 µg POPC measured from 8 different angles (light blue) and the average spectrum (blue).

Supplementary Fig. S3

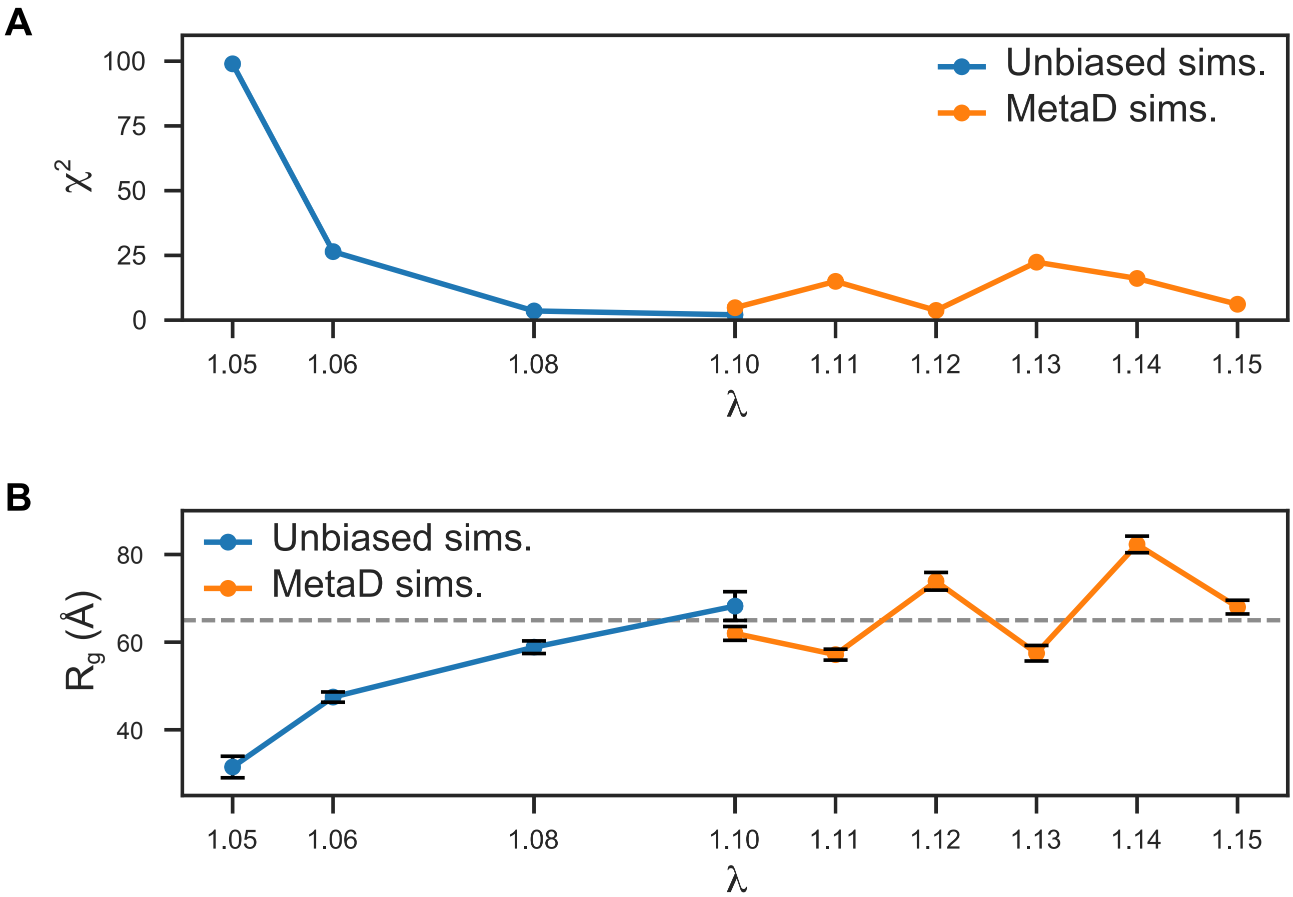


**Suppl. Fig. S3: Calibration of the protein-water interactions strength on the Martini3 forcefield for the simulation of hGHR-ICD.** The strength of the protein-water interactions was varied by a factor (λ) in the range between 5 and 10% in unbiased MD simulations (5%: 5 µs; 6%: 3 µs; 8%: 3µs and 10%: 5µs) and 10%-15% in metadynamics simulations (all of them ~ 10 µs) to improve the sampling. Backmapped (CG→AA) conformations (1/ns) from each simulation were used to fit the SAXS data of hGHR-ICD 1.1 mg/mL as described in the *Materials and Methods* section. (A) Average $\chi^2$ obtained from fitting to the SAXS data and (B) Average $R_g$ measured from the conformations taken from the unbiased (blue) and metadynamics (orange) simulations. The dashed gray line in B corresponds to the hGHR-ICD $R_g$ obtained from the SAXS data.

Supplementary Fig. S4

**A**

| Protein | POPC lipid |
|---|---|
| Monomer: hGHR(MSP1D1) (F3) | 122 ± 17 |
| Dimer: hGHR(MSP1D1) (F1) | 115 ± 19 |
| MSP1D1* | 120 ± 10 |

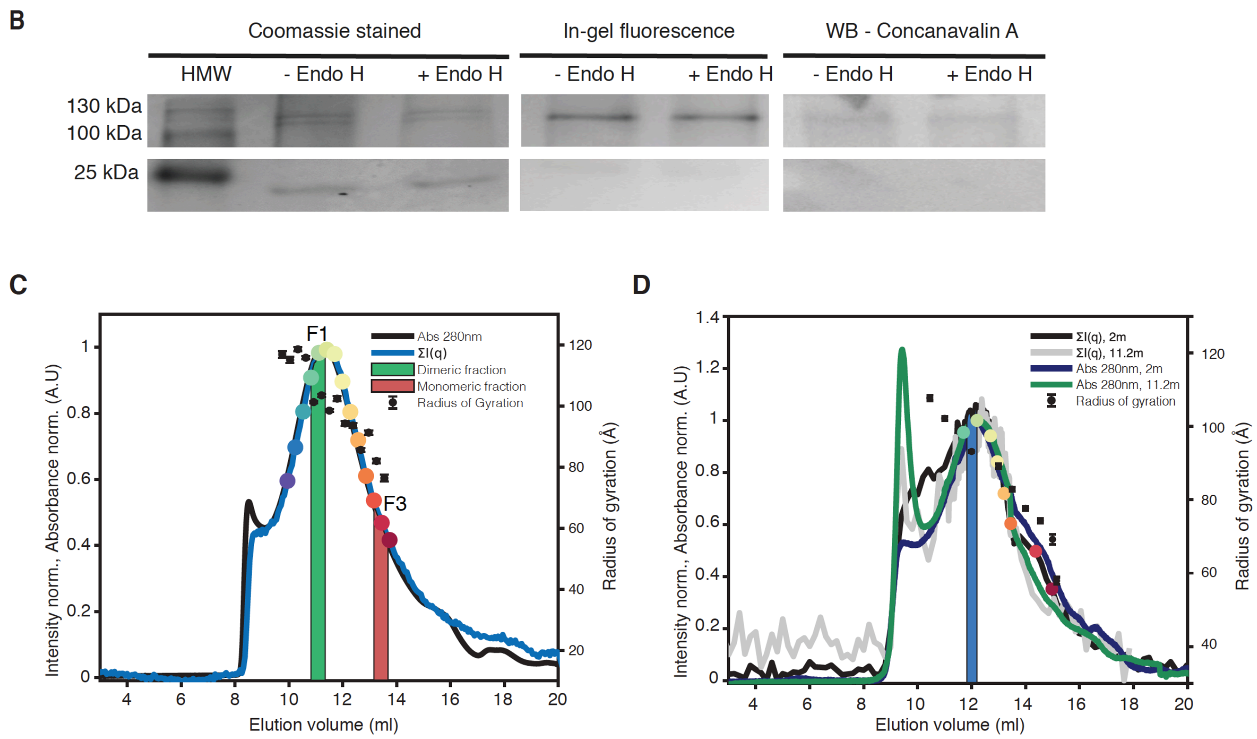


**Suppl. Fig. S4: Biochemical and biophysical analyses of hGHR in POPC loaded nanodiscs**. (A) POPC/hGHR-loaded MSP1D1 ratios were derived from fraction 1 (F1, mostly dimer) and 3 (F3, monomer) by phosphate analysis (see *Materials and Methods*). (B) hGHR-loaded MSP1D1 was treated with Endo-H overnight (see *Materials and Methods*). Endo-H (+Endo-H) and untreated (-Endo-H) were separated by SDS-PAGE and analysed by in-gel fluorescence, Coomassie staining, and western blotting using horse-radish peroxidase conjugated Concanavalin A that binds mannose residues. (C) SEC-SAXS data for hGHR in MSP1D1 with POPC. Absorption at 280 nm (black line), scaled to the total scattering intensities of individual frames (blue line). Calculated $R_g$ values are plotted as black dots. The green and red areas indicate the fractions chosen for the dimer and the monomer respectively. The SAXS data corresponding with the green and red areas are plotted in Figure 4F. (D) SEC frames corresponding to the two SEC-SANS data sets, one for each detector setting. Absorption at 280 nm (navy and green lines), scaled to the total scattering intensities obtained for the two settings, (black and grey lines). Calculated $R_g$ values are plotted as black circles. The SANS data corresponding with the blue area is plotted in Fig. 4G.

Supplementary Fig. S5

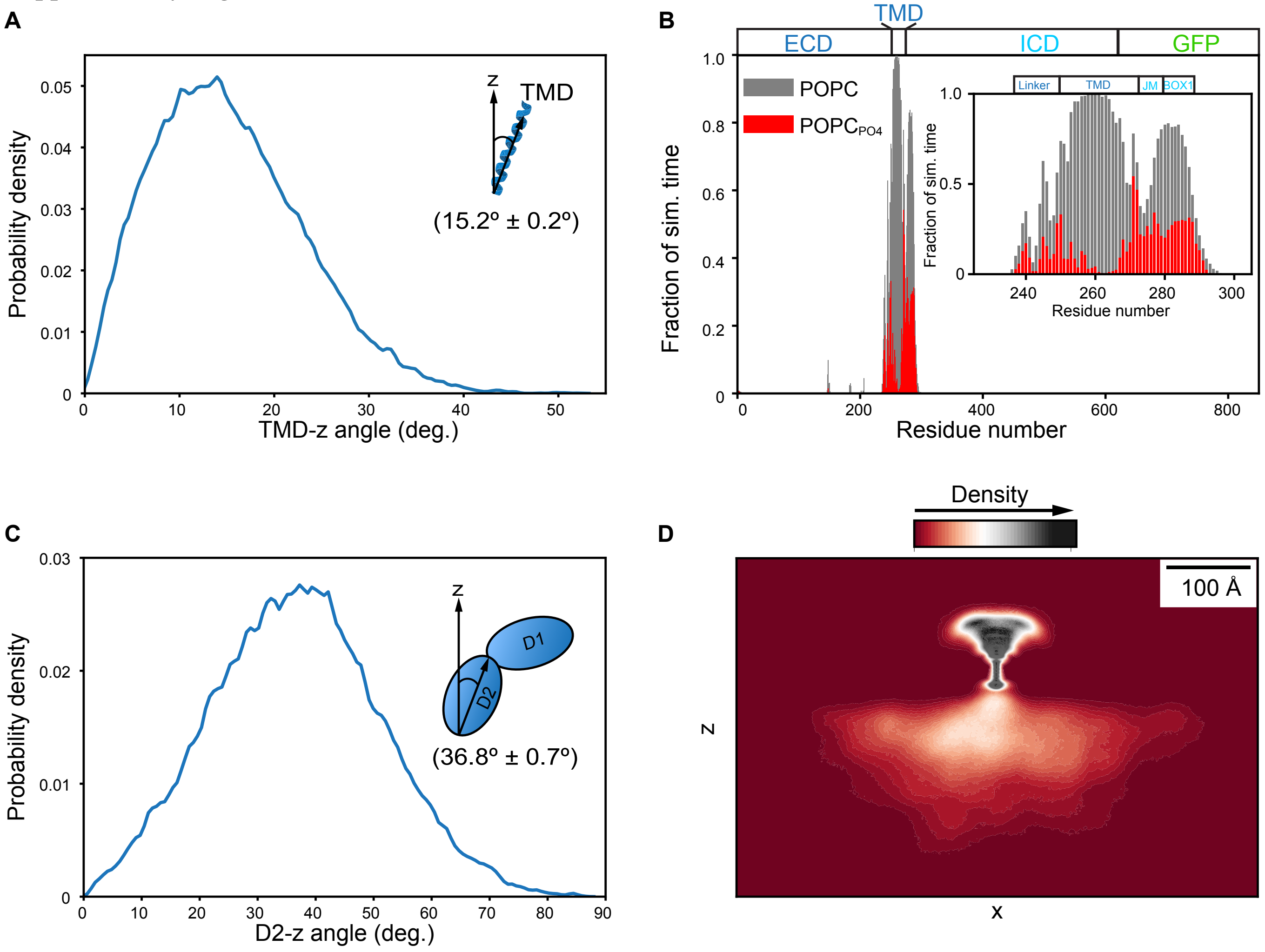


**Suppl. Fig. 5. Additional insights of hGHR-GFP relative orientations from the CG-MD simulation of hGHR-GFP in a POPC bilayer.** (A) Probability density of the angle between the principal axis of the TMD and the z-axis (perpendicular to the membrane plane). (B) Protein-lipid contact profiles. A contact is defined when the distance between a BB bead of the protein and a $PO_4$ bead (red) or any bead (gray) of POPC is ≤ 7 Å. The inset shows a detail of the profile highlighting the interactions between the intracellular juxtamembrane (ICJM) and BOX1 regions of GHR-ICD with the lipids. (C) Probability density of the angle between the principal axis of the D1 domain and the z-axis. (D) Average volumetric protein (hGHR-GFP) density map projected on the *xz* plane. This map provides a visualization of the large volume that the ICD+GFP can occupy below the membrane in contrast with the ECD. In all panels the measurements were performed considering the last 20 µs of the hGHR-GFP+$POPC_{pws10}$ simulation, hence the first µs was discarded as equilibration time.

**Supplementary Tables**

**Suppl. Table S1.** Molecular weights of the hGHR-ECD and hGHR-ICD from the SAXS data.

| Protein sample | *I(0)* (1/(mg/ml)) | Concentration (mg/mL) | Theoretical molecular weight (kDa) | Calculated molecular mass (kDa) |
|---|---|---|---|---|
| hGHR-ECD | 0.083 | 3.45 | 28.1 | 33.3 |
| hGH | 0.032 | 1.83 | 24.2 | 22.1 |
| hGH:hGHR-ECD1:2 | 0.067 | 1.28 | 72.5 | 78.2 |
| hGH:hGHR-ECD1:1 | 0.012 | 0.30 | 55.4 | 50.2 |
| hGHR-ICD | 0.037 | 1.14 | 38.6 | 45.0 |
| hGHR-ICD-GFP-$H_{10}$ | 0.05 | 0.90 | 68.0 | 75.8 |

**Suppl. Table S2.** Dimensions of hGHR-ICD (1.14 mg/ml sample) – Comparisons of different analysis approaches

| **Analysis** | $R_g$ (Å) | $R_h$ (Å) | ν | Reference |
|---|---|---|---|---|
| **SAXS** | | | | |
| Guinier | 64.5± 1.3 | | **n.a.** | |
| IFT | 62.0 ± 0.1 | | **n.a.** | [3] |
| GRC model fit | 68 ± 4 | | 0.5** | [4] |
| Extended Guinier | 65.9 | | 0.618±0.002 | [5] |
| Sosnick | 64.9 ± 0.4 | | 0.603±0.002 | [6] |
| Kohn* | 64.2 | | 0.598** | [7] |
| **NMR** | | | | |
| Diffusion NMR | | 43.9±0.5 | **n.a.** | |
| Forman-Kay* | | 49.2 | 0.509** | [8] |

*: Empirical prediction based on the number of residues of hGHR-ICD (352).
**: Parameter prefixed in the model.
SAXS analysis is based on the hGHR-ICD sample of 1.14 mg/ml (see main text).
See main text and Materials and Methods for description of the NMR conditions.

**Suppl. Table S3:** SAXS-refined fit parameters of the hGHR-GFP in nanodisc

| **Fitting Parameters** | |
|---|---|
| $A_{head}$ (Å$^2$) | 66 ± 10 |
| $R_g$ of coil (Å) | 76 ± 8 |
| $\nu_{POPC}$ (Å$^3$) | 1230 ± 10 |
| $\nu_{GHR-GFP}$ (Å$^3$) | 119 600 ± 3400 |
| $H_{belt}$ (Å) * | 25.8 |
| ε * | 1.5 |
| $N_{POPC}$ * | 122 |
| Roughness (Å) * | 5.0 |
| **Deduced parameters** | |
| $H_{ND}$ (Å) | 37.7 |
| $D_{minor}$ (Å) | 29.6 |
| $D_{major}$ (Å) | 44.3 |
| $d_{belt}$ (Å) | 8.16 |

*Parameter not fitted.

$A_{head}$: area taken up by one POPC headgroup.

$R_g$ of coil: ensemble average $R_g$ of the Gaussian random coil representing the ICD.

$\nu_{POPC}$: partial specific molecular volume of one POPC molecule.

$\nu_{GHR-GFP}$: partial specific volume of the hGHR-GFP.

$H_{belt}$: height of the MSP-belt.

ε: axis ratio of the elliptical phospholipid bilayer patch ($D_{major}$ / $D_{minor}$) .

$N_{POPC}$: the average number of POPC per nanodisc.

Roughness: Interface rougness correcting for the fact the interfaces are not perfectly smooth.

$H_{ND}$: total height of the phospholipid bilayer.

$D_{minor}$ and $D_{major}$: major and minor diameter of the phospholipid bilayer.

$D_{belt}$: thickness of the MSP belt.

**Supplementary text**

**Detailed account of the results of the semi-analytical model fit of hGHR-GFP in POPC nanodiscs**

To refine our model against the data from full-length GHR-GFP in nanodiscs using a semi-analytical model, a total of five parameters were fitted: the area per one POPC headgroup; the ensemble average $R_g$ of the Gaussian random coil representing the disordered intracellular domain; the partial specific molecular volumes of the POPC lipids ($\nu_{POPC}$) and the hGHR-GFP membrane protein ($\nu_{GHR\text{-}GFP}$), respectively and finally a small constant background in order to correct for small errors in the background subtraction. $\nu_{POPC}$ and $\nu_{GHR\text{-}GFP}$ were allowed to vary within a few percent of their pre-estimated values. The initial value for $\nu_{POPC}$ was taken from literature as 1260 Å$^3$,[9] whereas the initial partial specific molecular volumes of one MSP and the ICD of the GHR, $\nu_{MSP}$ and $\nu_{GHR\text{-}ICD}$, were calculated using an average mass density of proteins of 1.35 cm$^2$/g to be 27 100 Å$^3$ and 47 300 Å$^3$, respectively. The partial specific molecular volumes of the remaining part of the GHR, *i.e.* the GHR-ECD-TMD and the GFP were calculated as part of *WillItFit*[10–12], giving a total volume of $\nu_{GHR\text{-}GFP}$=116 600Å$^3$ for the full-length GHR-GFP.

In the upper part of supplementary Table S3, the parameter values of the five main parameters are listed along with the fixed model parameters. In the lower part of the table, parameters deduced from the fit parameters due to the systematic use of molecular constraints are listed. Aside from the listed parameters, a small constant value was added to the model in order to correct for errors in the background subtraction.

In the fits, the belt-height was fixed to 25.8 following the same arguments as previously outlined[13]. The value for the interface roughness was fixed to 5Å and the value for the axis ratio of the elliptical cylinder representing the phospholipid bilayer was fixed to 1.5, both in line with the results from refinements of the nanodisc model on comparable nanodisc systems[10,12,13].

### SUPPLEMENTARY REFERENCES


1. Wei, Y., Thyparambil, A. A. & Latour, R. A. Protein helical structure determination using CD spectroscopy for solutions with strong background absorbance from 190 to 230nm. *Biochim. Biophys. Acta - Proteins Proteomics* **1844**, 2331–2337 (2014).
2. Sommese, R. F., Sivaramakrishnan, S., Baldwin, R. L. & Spudich, J. A. Helicity of short E-

R/K peptides. *Protein Sci.* **19**, 2001–2005 (2010).
3. Hansen, S. BayesApp: A web site for indirect transformation of small-angle scattering data. *J. Appl. Crystallogr.* **45**, 566–567 (2012).
4. Debye, P. Molecular-weight determination by light scattering. *J. Phys. Colloid Chem.* **51**, 18–32 (1947).
5. Zheng, W. & Best, R. B. An Extended Guinier Analysis for Intrinsically Disordered Proteins. *J. Mol. Biol.* **430**, 2540–2553 (2018).
6. Riback, J. A. *et al.* Innovative scattering analysis shows that hydrophobic disordered proteins are expanded in water. *Science* **358**, 238–241 (2017).
7. Kohn, J. E. *et al.* Random-coil behavior and the dimensions of chemically unfolded proteins. *Proc. Natl. Acad. Sci. U. S. A.* **101**, 12491–6 (2004).
8. Marsh, J. A. & Forman-kay, J. D. Sequence Determinants of Compaction in Intrinsically Disordered Proteins. *Biophysj* **98**, 2383–2390 (2010).
9. Kucerka, N., Tristram-Nagle, S. & Nagle, J. F. Structure of fully hydrated fluid phase lipid bilayers with monounsaturated chains. *J. Membr. Biol.* **208**, 193–202 (2005).
10. Skar-Gislinge, N. *et al.* Small-angle scattering determination of the shape and localization of human cytochrome P450 embedded in a phospholipid nanodisc environment. *Acta Crystallogr. D. Biol. Crystallogr.* **71**, 2412–21 (2015).
11. Pedersen, M. C., Arleth, L. & Mortensen, K. WillItFit : a framework for fitting of constrained models to small-angle scattering data. *J. Appl. Crystallogr.* **46**, 1894–1898 (2013).
12. Kynde, S. A. R. *et al.* Small-angle scattering gives direct structural information about a membrane protein inside a lipid environment. *Acta Crystallogr. Sect. D Biol. Crystallogr.* **70**, 371–383 (2014).
13. Johansen, N. T. *et al.* Circularized and solubility-enhanced MSPs facilitate simple and high-yield production of stable nanodiscs for studies of membrane proteins in solution. *FEBS J.* **286**, 1734–1751 (2019).
